# Evaluation and Optimization of High-Field Asymmetric Waveform Ion Mobility Spectrometry for Multiplexed Quantitative Site-specific N-glycoproteomics

**DOI:** 10.1101/2021.03.23.436434

**Authors:** Pan Fang, Yanlong Ji, Ivan Silbern, Rosa Viner, Thomas Oellerich, Kuan-Ting Pan, Henning Urlaub

## Abstract

The heterogeneity and complexity of glycosylation hinder the depth of site-specific glycoproteomics analysis. High-field asymmetric-waveform ion-mobility spectrometry (FAIMS) has shown to improve the scope of bottom-up proteomics. The benefits of FAIMS for quantitative N-glycoproteomics have not been investigated yet. In this work, we optimized FAIMS settings for N-glycopeptide identification, with or without the tandem mass tag (TMT) label. The optimized FAIMS approach significantly increased the identification of site-specific N-glycopeptides derived from the purified IgM protein or human lymphoma cells. We explored in detail the changes in FAIMS mobility caused by N-glycopeptides with different characteristics, including TMT labeling, charge state, glycan type, peptide sequence, glycan size and precursor m/z. Importantly, FAIMS also improved multiplexed N-glycopeptide quantification, both with the standard MS2 acquisition method and with our recently developed Glyco-SPS-MS3 method. The combination of FAIMS and Glyco-SPS-MS3 provided the highest quantitative accuracy and precision. Our results demonstrate the advantages of FAIMS for improved mass-spectrometry-based qualitative and quantitative N-glycoproteomics.

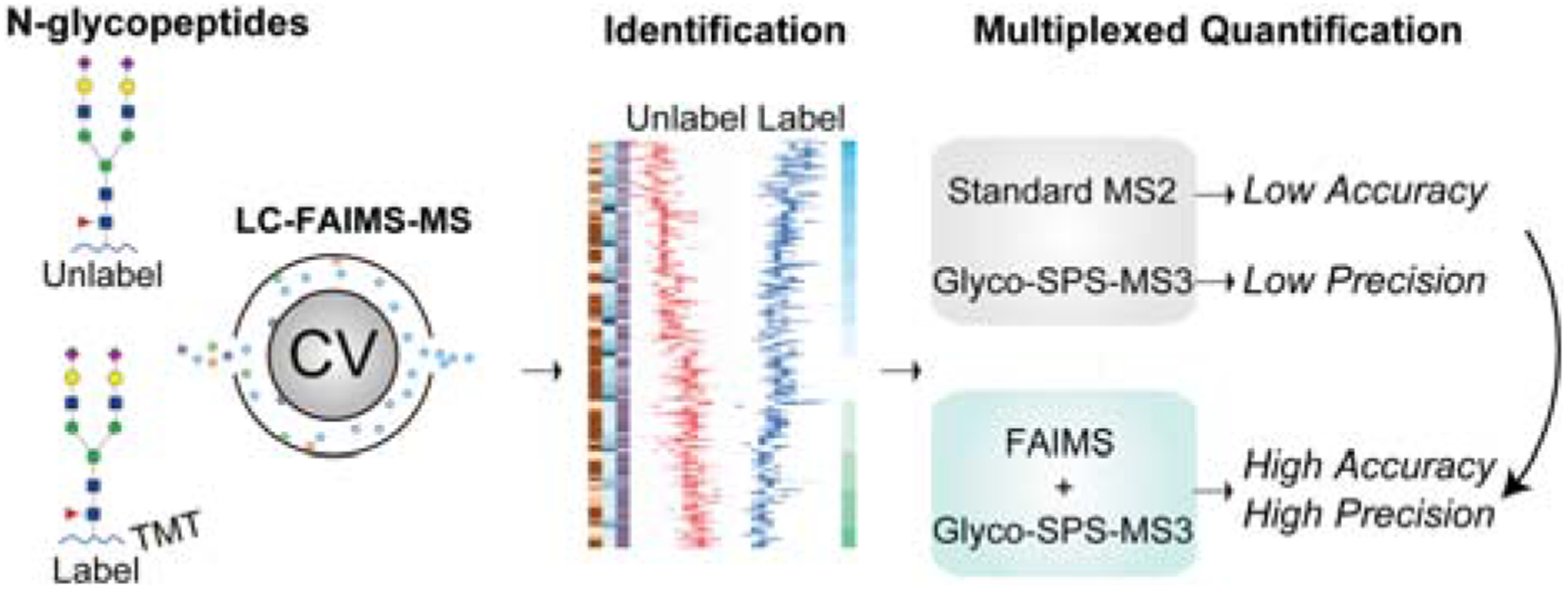

## INTRODUCTION

N-glycosylation is one of the most prominent and abundant post-translational modifications of proteins and plays fundamental roles in various biological processes.^1^ In-depth identification and quantification of the N-glycoproteome in physiological and pathological conditions are critical for an understanding of the functional roles of N-glycosylated proteins.^2, 3^ Continuous development in mass spectrometry (MS)-based N-glycoproteomics have made large-scale global analysis possible, with more confident N-glycopeptide identification, using streamlined workflows. ^4^ However, owing to the structural diversity of glycans and the correspondingly heterogeneous nature of N-glycopeptides, the overall sensitivity of N-glycoproteomics still requires further improvement.

One especial challenge in N-glycoproteomics is deciphering the micro-heterogeneity of individual glycosylation sites in a comprehensive manner. In C18 reversed-phase liquid chromatography (LC)-MS, N-glycopeptides sharing a unique peptide sequence but carrying different glycans (i.e., micro-heterogeneous glycopeptides) are detected at closely spaced retention times with considerable intensity differences. These often demand a greater number of accumulated ions for MS acquisition, and consequently longer MS scan time, if informative fragment ions of N-glycopeptides are to be identified reliably.^5, 6^ This makes it very difficult for a standard LC-MS/MS analysis to capture all the glycopeptides in a complex sample, in the time available.

Continuous developments in ion mobility-mass spectrometry, which have made the addition of gas-phase separation to an LC-MS/MS analysis possible, enhanced analyte separation and characterization.^7^ Among different ion mobility technologies, high-field asymmetric-waveform ion-mobility spectrometry (FAIMS) reportedly improves the range and reliability of bottom-up proteomics and phosphoproteomics.^8–18^ FAIMS separates ions on the basis of their differential gas-phase mobility in alternating low and high electric fields between inner and outer electrodes. The mobility of an ion depends on several properties, including its mass, shape, center of mass, dipole moment and neutral-gas– ion interaction.^19^ As a result, FAIMS is orthogonal to LC and can be integrated into an LC-MS system (i.e., LC-FAIMS-MS). A DC potential, termed the compensation voltage (CV), applied to the FAIMS electrode allows selective transmission of ions through the device before they enter the mass spectrometer.

In addition to proteomics and phosphoproteomics, where different CVs in an LC-FAIMS-MS measurement have been applied to cover complementary gas-phase fractions, LC-FAIMS-MS has also shown increased power to identify cross-linked peptides and bacterial glycopeptides.^20, 21^ Since protein glycosylation in human cells differs significantly from that in bacteria, it is necessary to adapt empirically the optimum CV settings for human N-glycoproteome. Especially, it remains unclear how the various glycan structures influence the FAIMS separation of glycopeptides.

Multiplexed quantitative N-glycoproteomics enhances sample throughput while maintaining quantification performance.^2, 22^ However, quantification based on isobaric chemical labeling is limited by the co-isolation and co-fragmentation of interfering ions within the target precursor isolation window, resulting in distorted reporter-ion ratios.^23^ We have recently extended the synchronous precursor selection (SPS)-MS3 method^24^ and developed Glyco-SPS-MS3^5^ to improve the quantification accuracy for multiplexed N-glycoproteomics. Schweppe et al. have demonstrated the ability of FAIMS to remove background ions and improve quantification accuracy in multiplexed proteomics using MS2 and MS3 methods.^12^

In this work, we aimed to systematically evaluate different FAIMS settings for the quantification of tandem mass tag (TMT)-labeled heterogeneous N-glycopeptides. Since chemical labeling can affect FAIMS settings, we comprehensively assess the changes in FAIMS-aided separation of N-glycopeptides with and without labeling with the TMT. The optimum FAIMS settings performed just as well with N-glycopeptides derived from purified IgM as they did with N-glycopeptides from DG75 cells. In detail, we investigated the effects of different N-glycopeptides’ characteristics on FAIMS separation, including TMT labeling, charge state, glycan type, peptide sequence, glycan size and precursor m/z. Finally, using an IgM-yeast model, we compared the abilities of the MS2 and MS3 methods, operating with and without FAIMS, to perform accurate and precise multiplexed quantification of N-glycopeptides.

## EXPERIMENTAL SECTION

### Sample Preparation

Human IgM purified from human serum (Sigma) was digested with trypsin, and the resulting peptides were labeled without or with TMT reagents (TMT0 or TMT6). The IgM digests (without enrichment of glycopeptides) were injected onto LC-FAIMS-MS/MS analysis directly. DG 75 cells (DSMZ no.: ACC 83) were digested with trypsin and the resulting peptides were then labeled with TMT tag (TMT0 or TMT6) followed by enrichment of glycopeptides using zwitterionic hydrophilic interaction liquid chromatography (ZIC-HILIC). IgM and yeast peptide mixture were prepared as showed in **Fig. 5A**. The detailed experiments for all samples were supplemented in **Supporting Information**.

### LC-FAIMS-MS/MS Analysis

For LC-MS/MS measurements with FAIMS, the Thermo FAIMS Pro device (Thermo Fisher Scientific) was operated in standard resolution with the temperature of FAIMS inner and outer electrodes set to 100 ^o^C. The dispersion voltage (DV) circuitry was tuned using the auto-tune option. The option “Tune DV RF” was enabled throughout the LC-FAIMS-MS/MS analysis for the high and low electric field’s automated settings that create the DV waveform applied to the electrodes. In order to maintain a stable electrospray, we used a coated silica emitter (New Objective) for LC-MS/MS runs with FAIMS, while a metal emitter (Thermo Fisher Scientific) was used for no FAIMS analyses. For single-CV experiments, a CV ranging from −25 to −90 V with 5 V steps was applied throughout the analysis. For runs with double CVs or triple CVs, selected CV was applied to sequential survey scans and MS/MS cycles. The cycle time was 1.5 s for each CV in double CV experiments and 1 s for each CV in triple CV experiments. The MS/MS CV was always paired with the appropriate CV from the corresponding survey scan. LC-FAIMS-MS2/MS3 methods were performed on the Orbitrap Exploris and Orbitrap Fusion mass spectrometers (Thermo Fisher Scientific). For the FAIMS CV scanning of synthetic peptides and glycopeptides, each sample (**Table S1, Supporting Information**) was directly injected to the Orbitrap Lumos (Thermo Fisher Scientific, San Jose, CA). The CV was scanned from 0-100 V in 1 V steps. For each run, selected ion monitoring scan type was used to monitor the ions with specific mass to charge (m/z). The parameters for LC and MS analyses were provided in detail in **Supporting Information**.

### Data Analysis

For intact N-glycopeptide identification and quantification,.raw files were processed via GlycoBinder.^5^ Intensities of all isotopic clusters associated with the identified N-glycopeptide were determined by the pQuant algorithm incorporated in pGlyco2.0.^25^ MaxQuant was used to determine the charge state and m/z of MS1 features in LC-MS/MS data.^26^ Parameters used for database search for IgM, DG 75 and IgM-yeast mixture samples were detailed in **Supporting Information**.

To assess the quantitative accuracy of different LC-FAIMS-MS methods, we first calculated the error of each TMT-ratio determination compared to the predefined values. We then deduced the “positive bias” of each acquisition method by taking the mean of all positive values of the errors associated with the method. For assessing quantitative precision, we calculated the population variance of each measurement based on the formula: population 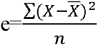, where X is the determined TMT ratio, 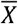 is the mean of all TMT ratios, and n is the count of all TMT ratios.

We classified all identified N-glycopeptides and their glycan compositions, i.e., numbers of different monosaccharide moieties including hexose (Hex), N-acetylhexosamine (HexNAc), N-acetylneuraminic acid (Neu5Ac) and Fucose (Fuc), into five putative glycan types based on previous glycomics analysis.^27, 28^ The potential number of branches of each glycan composition were proposed based on the glycan types. The detailed criteria for glycan classification were supplemented in **Supporting Information** and **Table S2**.

### Data Availability

The mass spectrometry raw data and related research results have been deposited to the ProteomeXchange Consortium via the PRIDE partner repository with the dataset identifier **PXD 023951**.^29^.

## RESULTS AND DISCUSSION

### Optimization of FAIMS CV settings for identification of TMT-labeled and unlabeled N-glycopeptides

To assess the effects of FAIMS CV settings on N-glycopeptide identification, we analyzed TMT-labeled and unlabeled human IgM tryptic digests using different single CVs, ranging from −25 to −90 V (in 5 V steps) in LC-FAIMS-MS/MS analyses. The resulting raw MS data were processed for N-glycopeptide identification by using the pGlyco 2 algorithm.^6^ As shown in **Figure 1A**, we identified the most TMT-labeled unique IgM N-glycopeptides with CVs ranging from −45 to −50 V, while obtaining the highest number of N-glycopeptide-spectra matches (GPSMs) at a CV of −55 V. Optimum CVs for unlabeled N-glycopeptides showed different settings. A broader range of CVs, from −45 to −65 V, performed similarly well for unique identifications of unlabeled IgM N-glycopeptides, whereby the CVs of −60 and −65 V led to the highest number of GPSMs (**Figure 1B**). Since FAIMS selects ions and diverts unwanted ions from entering the mass spectrometer, the application of only single CVs resulted in fewer unique N-glycopeptide identifications than without FAIMS. Nonetheless, the optimum single CVs still led to a comparable number of unique N-glycopeptide identifications (in average 168 with FAIMS vs. 165 without FAIMS in TMT labeled samples and 87 vs. 96 in unlabeled samples) and even more GPSMs (in average 887 with FAIMS vs. 764 without FAIMS in TMT labeled samples and 345 vs. 301 in unlabeled samples), demonstrating an improved reliability in N-glycopeptide identification. To examine whether FAIMS improves the ion transmission of N-glycopeptides, we compared the ratios of glycan-oxonium-ion-containing spectra counts with the counts from all MS/MS spectra in each measurement across the range of CVs tested. Indeed, FAIMS increased the ratios of glycan-oxonium-ion-containing spectra by up to 65.5% (at CV −45 V) and 76% (at CV −50 V) for TMT-labeled and unlabeled samples, respectively (**Figure S1A**), enhancing the chance of N-glycopeptides being selected for fragmentation in an LC-MS/MS run. The difference in the optimum CVs for TMT-labeled and unlabeled N-glycopeptides (see above and compare **Figure 1A and 1B**) suggests that the attachment of TMT label influences the gas-phase mobility of N-glycopeptides in FAIMS. This means that the TMT-labeled N-glycopeptides possess properties different from those of unlabeled N-glycopeptides across tested CVs. Therefore, it was necessary to optimize the FAIMS CVs separately and specifically for N-glycopeptides with or without TMT-labeling.

**Figure 1.**
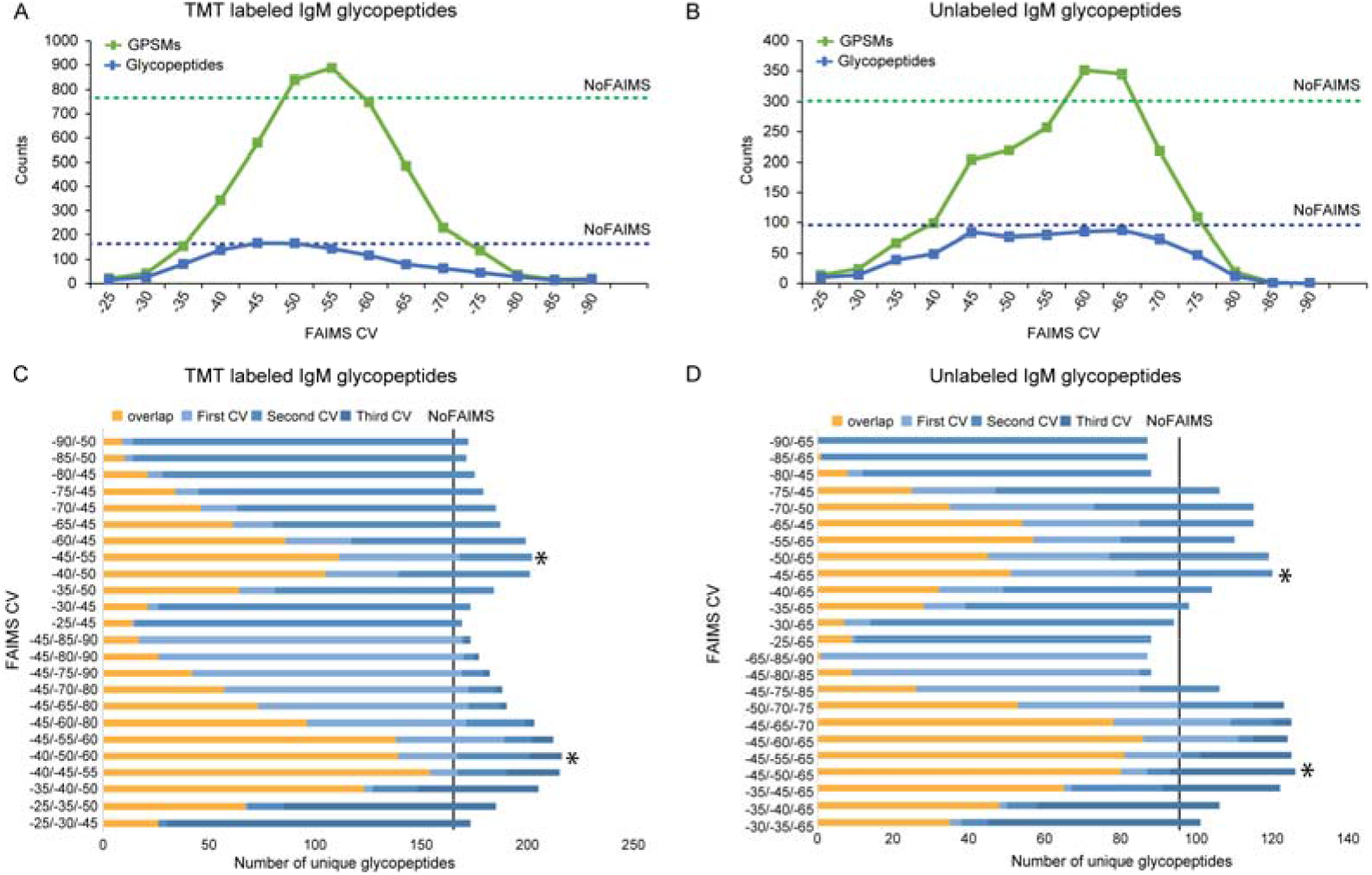
Evaluation and optimization of FAIMS CV settings for TMT-labeled and unlabeled IgM N-glycopeptides identification. (A, B) The numbers of identified GPSMs (green) and unique N-glycopeptides (blue) from IgM digests, (A) with and (B) without TMT labeling in LC-MS/MS analyses, using varying single CVs. The numbers of identified GPSMs and N-glycopeptides without FAIMS are shown as dashed lines. (C, D) The combined glycopeptide identifications of two and three CVs from IgM digests, (C) with and (D) without TMT labeling in LC-MS/MS analyses. The overlap and unique identifications of two-CV and three-CV settings are represented in yellow and blue, respectively. The optimal combination for two-CV and three-CV settings are indicated with asterisks. The numbers of identified unique N-glycopeptides without FAIMS are shown as vertical lines.

Earlier studies have shown that different CVs favor the identification of complementary groups of peptides with distinct physicochemical properties.^10^ Consequently, the combination of several CVs in one LC-FAIMS-MS analysis increases the overall coverage of peptide identification. We compared the identification of unique N-glycopeptides among any two CV settings and found that application of two close CVs led to up to 70% overlap (similarity) in detection and fragmentation (**Figure S1B**). We then determined the best CV setting-pairs for N-glycopeptide identification by combining the identifications of single CVs with a voltage range of −25 to −90 V in 5 V steps. For TMT-labeled glycopeptides with a two-CV combination, −45/-55 V provided the greatest number of identifications (**Figure 1C**). For unlabeled N-glycopeptides, −45/-65 V was found to be best (**Figure 1D**). We extended the analysis for three combined CV settings and found that the optimum CV settings were −40/-50/-60 V and −45/-50/-65 V for TMT-labeled and unlabeled N-glycopeptides, respectively. Application of the optimum two-CVs combination in a single analysis increased the overall N-glycopeptide identification/fragmentation by 20% for TMT-labeled glycopeptides and 25% for unlabeled glycopeptides. The optimized three-CV combinations allowed the identification of up to 30% more TMT-labeled and unlabeled N-glycopeptides as compared with no FAIMS measurement.

Next, we applied our CV evaluation based on IgM N-glycopeptide analyses to TMT-labeled N-glycopeptides derived from human DG75 cells. The CV of −55 V resulted in the largest number of identifications for unique N-glycopeptides with their respective GPSMs, and at −50 V for the highest numbers of N-glycosites, N-glycans and N-glycoproteins (**Figure 2A**). This observation agrees with those optimum CVs established with IgM-derived N-glycopeptides (**Figure 1** and above text). Since the DG75 sample is much more complex and richer in N-glycopeptides than the IgM sample, we also assessed the extent to which the combination of the different FAIMS CV settings and prolonged LC analysis time improves the depth of N-glycoproteomics in a single LC-FAIMS-MS analysis. To do this, we chose three LC gradients of different separation times (1, 2 and 3 h) with double-CV and triple-CV settings (**Figure 2B**). Analyses with prolonged LC separation time provided more identifications (unique N-glycopeptides and N-glycosites) using both double-CV and triple-CV settings. Compared with analyses with 1 h, separation with 2 h increased the overall number of N-glycopeptide identifications by 31% and 33% when double and triple CVs were applied, respectively. Separation with 3 h further improved the N-glycopeptide identification by 39% for double CV and 41% for triple CV. Application of triple-CV settings provided higher numbers of identifications at any separation time than that of double-CV settings. Compared with analyses without FAIMS, the application of double CV and triple CV at 3 h separation time showed a significant improvement in the identification of unique DG75 N-glycopeptides by a 2.6-fold and a 3.6-fold increase, respectively, (**Figure 2C**).

**Figure 2.**
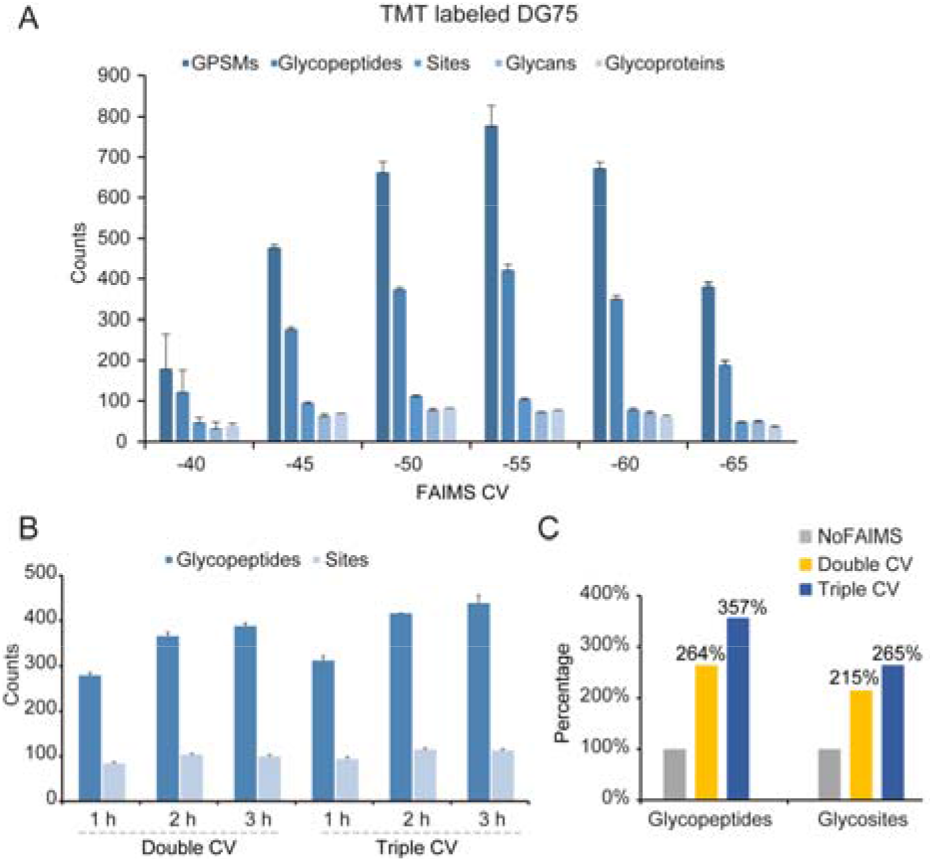
Evaluation and optimization of FAIMS CV settings for identifying TMT-labeled N-glycopeptides derived from DG75 cells. (A) The counts of identified GPSMs, unique N-glycopeptides, N-glycosites (Sites), glycan compositions and glycoproteins using single CVs ranging from −40 V to −65 V with steps of 5 V. (B) Evaluation of gradients (1 h, 2 h, 3 h) for FAIMS analysis using the double (−45/−55 V) and triple (−45/−50/−55 V) CVs. (C) Increase in the number of identified unique N-glycopeptides and N-glycosites from DG75 cells using double CV (−45/−65 V) and triple CV (−45/−60/−70 V), as compared to no FAIMS settings. Error bars represent the standard deviation from two FAIMS measurements.

### Characteristics of TMT-labeled and unlabeled IgM N-glycopeptides in LC-FAIMS-MS

In agreement with earlier studies,^10, 12, 20^ FAIMS significantly reduced the number of detectable singly charged precursors in LC-MS analyses (**Figure S2A**). The ratios of singly charged ions to all detectable ions in all MS1 features detected with CVs of −40 to −90 V were 50% less than that without FAIMS, for both TMT-labeled and unlabeled N-glycopeptides, revealing efficient suppression of singly charged background ions. The attachment of the TMT tag affected the charge state distribution of both MS1 features (**Figure S2A**) and identified N-glycopeptides (**Figure S2B**) across the CV settings evaluated. Interestingly, although TMT labeling led to differences in optimum CV settings for the overall N-glycopeptides identification by only 10 V difference (comparing **Figure 1A and 1B**), we noticed that differences in CV settings had a strong impact on the identification of individual N-glycopeptides (**Figure 3A**). We compared the MS1 intensities of all identified IgM N-glycopeptides with and without TMT labeling across the range of applied CVs and determined the CV settings that provided the most intense MS1 signal for each pair of unlabeled and labeled N-glycopeptides. We found that the differences between CVs (i.e., Δ CV) yielding the most intense MS1 signals from the same unlabeled and labeled glycopeptide can vary within a broad range, from −45 to +45 V (**Figure 3A**). N-glycopeptide ions with different charge states or strongly varying numbers of sialic acid (a negative-charge-bearing glycan moiety) showed distinct distributions of Δ CV values (**Figure S3**). This suggests that the TMT-induced changes in FAIMS mobility of some N-glycopeptides are largely charge-dependent. Indeed, triply charged N-glycopeptides were better identified at significantly higher CVs than did those that were quadruple charged (**Figure 3B**).

**Figure 3.**
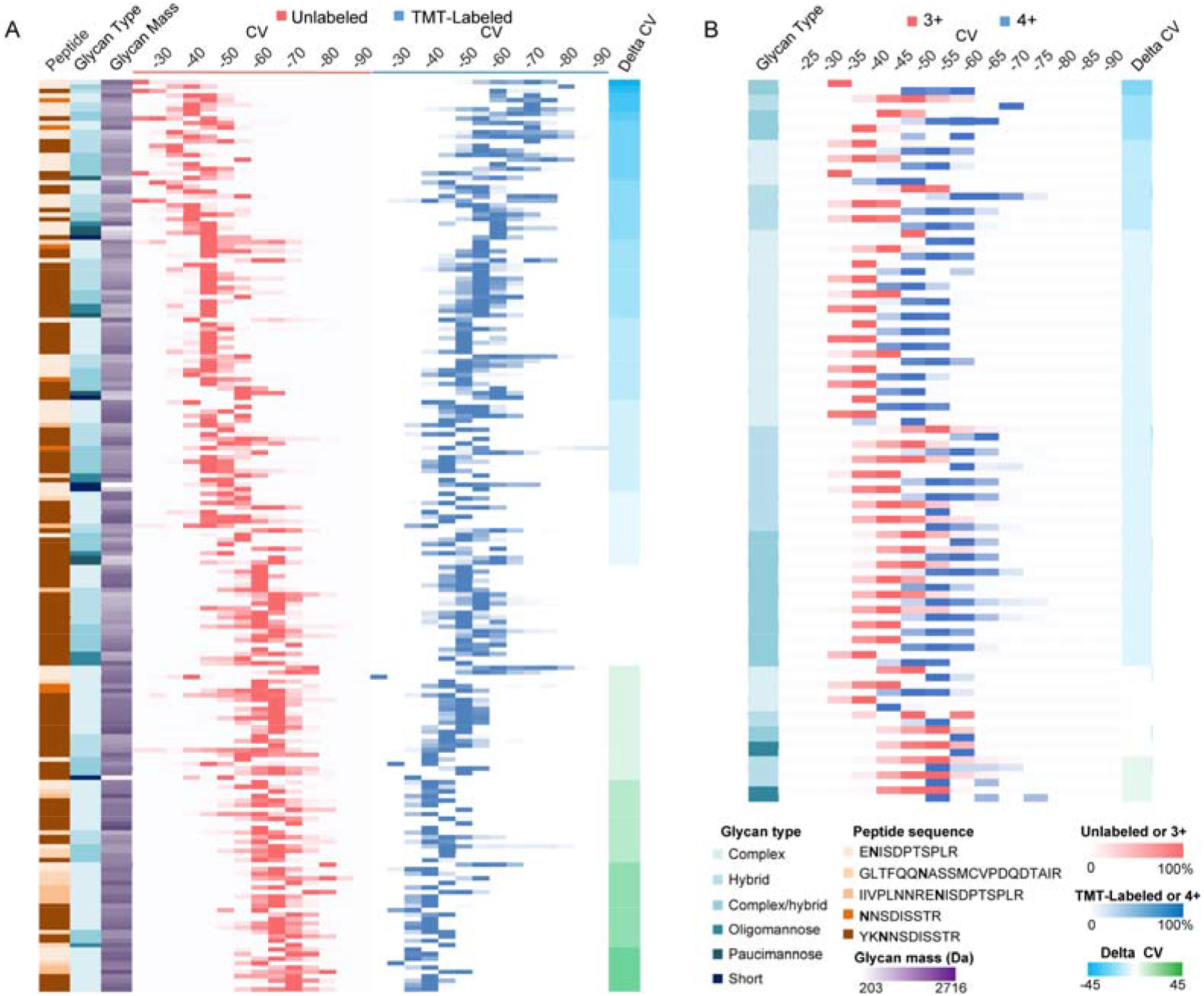
The influence of TMT labeling and charge states on the FAIMS mobility of IgM N-glycopeptides. (A) Normalized MS1 intensities of each pair of TMT-labeled (blue) and unlabeled (red) IgM N-glycopeptides detected with different CVs are shown on color scales. Δ CV is the difference obtained by subtraction of the optimum CV (the CV that provides the strongest intensity) of a TMT-labeled glycopeptide from that of the unlabeled counterpart. Characteristics of the N-glycopeptides and Δ CV values are color-coded, as shown in the figure. (B) Normalized MS1 intensities of each pair of IgM N-glycopeptides bearing the charges 3+ (red) and 4+ (blue) detected with different CVs are shown on color scales. Only N-glycopeptides containing the same IgM peptide sequence (YK<u>**N**</u>NSDISSTR) and modification type (TMT-labeled N-terminus and lysine) are considered.

To investigate more systematically the factors that influence the FAIMS mobility of N-glycopeptides bearing distinct glycans and peptide sequences, we prepared a series of synthetic N-glycopeptides and their non-glycosylated counterparts (**Table S1**) and injected them directly (i.e., via flow injection without LC separation) into an Orbitrap Lumos mass spectrometer for FAIMS CV plots. We monitored the MS1 intensities of each injected peptide/N-glycopeptide ion (m/z) while scanning FAIMS CVs from 0 to −100 V in 1 V steps (**Figure S4**). FAIMS separated the doubly and triply charged ions of peptides EVFVHPYSNK (P1) and EVFVHPYSDK (P2) but not EVFVHPYS<u>**N**</u>(GlcNAc)K (GP1). The 3+ and 4+ charge states of a larger bi-antenna sialyl-LacNAc glycan attached to EVFVHPYS<u>**N**</u>(5Hex4HexNAc2NeuAc) K (GP2) were separated as well. The charge states 2+, 3+, and 4+ of the longer glycopeptide YGNV<u>**N**</u>(5Hex4HexNAc2NeuAc) ETQNNSFK (GP4) were also separated efficiently by various FAIMS CV settings. Interestingly, for the same peptide but with 5Hex4HexNAc2NeuAc attached to C-terminal Asn YGNVNETQN<u>**N**</u>SFK (GP3), only charge state 3+ could be observed. In general, the optimum FAIMS CV for the four triply charged synthetic N-glycopeptides (GP1-4) differs only slightly and is independent of peptide length and glycan moiety. However, N-glycopeptides of different peptide length and sequence but carrying the same bi-antenna sialyl-LacNAc glycan (GP2 and GP4) showed a notable shift in optimum CV settings, especially for their 4+ charge states. Peptides sharing a common peptide sequence with or without the attachment of one monosaccharide moiety (P1 and GP1, doubly charged) were also readily separated by FAIMS. Overall, these initial results on synthetic N-glycopeptides suggest that their FAIMS mobility is influenced mainly by the charge state of the N-glycopeptide, glycan composition and the peptide sequence. To further disclose the impact of the glycan structure on optimum FAIMS CV settings, we determined the intensities of IgM-derived N-glycopeptides in LC-FAIMS-MS analyses with different CVs. Our results also revealed the influence of diverse glycan structures on optimum FAIMS settings. For instance, we found the while the optimum CVs for its individually glycosylated forms ranged from −25 to −75 V (**Figure S5**). optimum CV for the IgJ-derived non-glycosylated peptide E<u>**N**</u>ISDPTSPLR in the IgM sample to be −50 V,

To explore systematically the effect of different glycans on the N-glycopeptide identification in LC-FAIMS-MS analysis, we classified IgM N-glycopeptides into six groups according to their glycan compositions and compared the numbers of identified unique N-glycopeptides of these individual groups between different single CVs (**Figure 4A**). To exclude the effects of charge state, peptide sequence, and other modifications, we selected for evaluation only N-glycopeptides from human IgM with the same peptide sequence (YK<u>**N**</u>NSDISSTR), the same modification types (TMT6-modification at N-terminus and lysine residue for TMT labeled N-glycopeptides and no modifications for unlabeled N-glycopeptides) and the same charge state (3+). These were in total 100 unique N-glycopeptides for labeled IgM and 79 for unlabeled IgM. We found that the optimum single CVs for complex-type N-glycopeptides (−45 V for unlabeled N-glycopeptides and −40 V for labeled N-glycopeptides) were in general higher than those for hybrid types (−50 or −55 V for unlabeled N-glycopeptides and −45 V for labeled N-glycopeptides). N-glycopeptides bearing paucimannose or short glycans are better identifiable even at lower CVs (−75 V for unlabeled N-glycopeptides and −60 V for labeled N-glycopeptides). These data again confirmed that N-glycopeptides with increased numbers of negatively charged sialic acid moieties (complex and hybrid types) attached revealed optimum CV settings at elevated voltages (**Figure 4B**).

**Figure 4.**
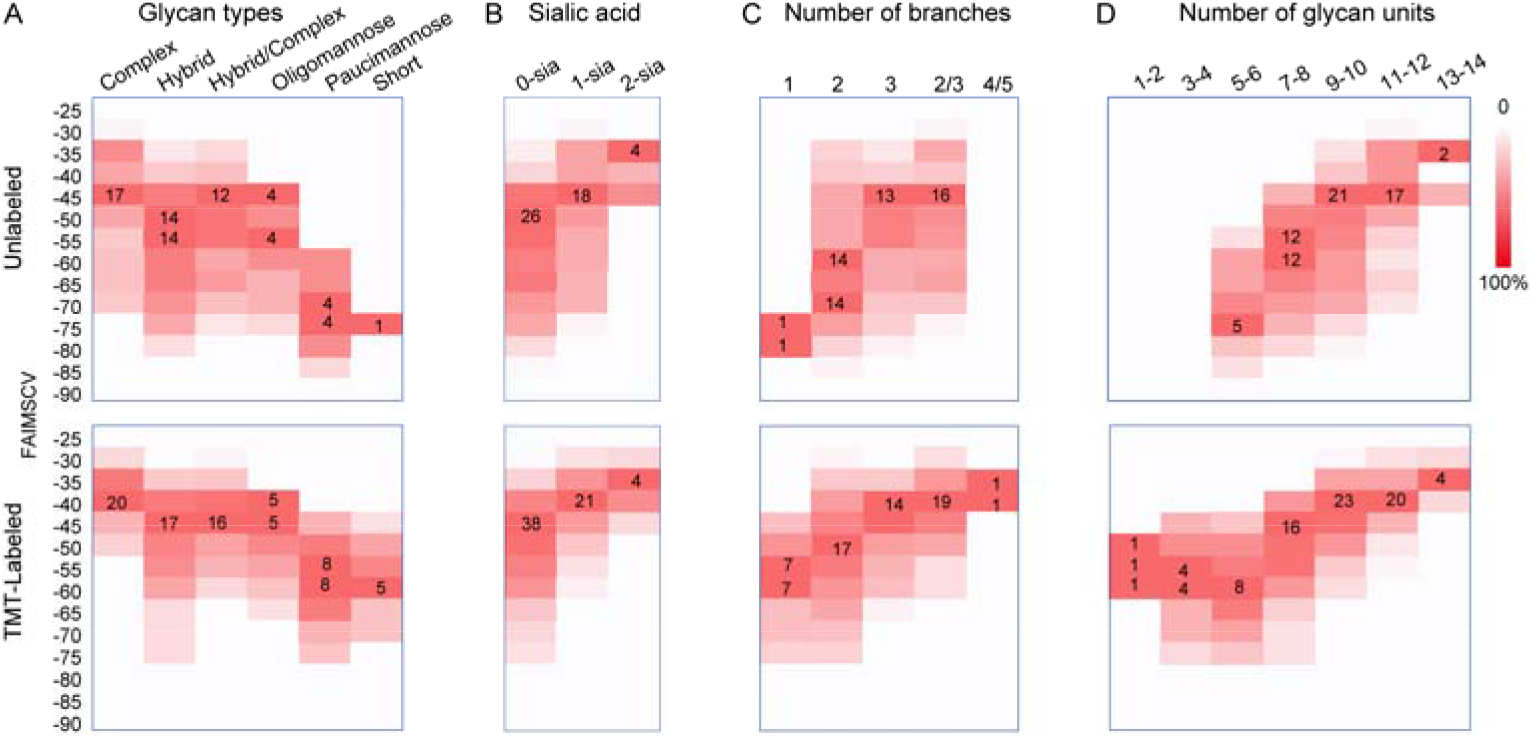
The influence of glycan structure and composition on the FAIMS mobility of TMT-labeled and unlabeled IgM N-glycopeptides. To preclude the influence of peptide sequence, modifications and charge state, we considered only the triply charged IgM N-glycopeptides carrying the peptide sequence “ YK<u>**N**</u>NSDISSTR” (TMT6-labeled N-terminal and lysine residue on the TMT-labeled N-glycopeptides and no modification on the unlabeled glycopeptides). We then classified the identified N-glycopeptides into different groups based on their glycan type (A), number of sialic acid moieties (B), number of possible branches (C), and number of glycan units (D). For each group, counts of unique N-glycopeptides identified with individual CVs were shown as a color scale in the figure. We also highlight the maximum unique N-glycopeptides counts of each group. The data represent the measurements of four replicates

The CVs that favor the identification of N-glycopeptides with individual/specific groups of glycans seemed to correlate with the size of the attached glycan. We thus calculated the number of monosaccharides in each identified glycan composition and proposed a potential number of branches based on classical human N-glycosylation pathway.^28^ Indeed, N-glycopeptides with attached glycans that have more potential branches or more sugar moieties required higher CVs (**Figure 4C, D**). The distributions of the precursor and glycan masses of all identified IgM and DG75 N-glycopeptides, in LC-MS/MS analyses with various single CVs, further confirmed such a correlation (**Figure S6 and S7**). In agreement with earlier FAIMS studies on proteomic, phosphoproteomic and protein crosslinking studies,^10, 17, 20^ our results show that the optimum CVs for N-glycopeptides strongly depend on the precursor m/z, and that N-glycopeptides with lower m/z showed better identifications at lower CVs.

Beside the CV-dependent ion-transmission efficiencies of FAIMS for N-glycopeptides with different glycan moieties, in-source fragmentation may also contribute to the detection of smaller or less heavily sialylated N-glycopeptides at lower CVs. Indeed, we found increased in-source fragmentation at lower CVs (lower than −55 V). For example, the N-glycopeptide “YK<u>**N**</u>NSDISSTR”, carrying a fucosylated bi-antenna N-glycan, was detected at a retention time (RT) of 16.5 min in a 60-min LC-MS/MS analysis, while the N-glycopeptide with one more sialic acid was often observed to elute later, at RT 20 min (**Figure S8**). However, we detected an increased signal from the non-sialylated form, alongside lowered intensities of the sialylated form, at RT 20 min when the CV was decreased from −45 to −60 V, indicating the occurrence of in-source fragmentation at lower CVs.

In summary, we showed that several factors can contribute to CV selection for FAIMS-assisted glycoproteomics, including TMT-labeling, charge state, glycan type, peptide sequence, glycan size and precursor m/z. These factors interplay and have mixed effects in influencing the mobility, and hence the detection and identification of N-glycopeptides. Amongst the various relevant factors, TMT labeling, charge state and precursor m/z showed the strongest influence on FAIMS mobility. While combining complementary CVs enhances the depth of glycoproteomics analysis, in-source fragmentation may occur at lower CVs and lead to the detection of non-natural N-glycopeptides.

### FAIMS improved accuracy and precision for multiplexed quantification of site-specific N-glycopeptides

Since FAIMS removes background ions and reduces the complexity of ions injected into the mass spectrometer, it may alleviate co-isolation interference in multiplexed quantitative glycoproteomics and thus improve quantitation of labeled N-glycopeptides. Very recently, we established a Glyco-SPS-MS3 method^5^ to reduce co-isolation interference and improve the quality of N-glycopeptide fragmentation and hence the overall accuracy of quantification. We therefore investigated whether the application of FAIMS was beneficial in enhancing quantitative accuracy in complex glycoproteomic samples. To benchmark the FAIMS-assisted quantitative N-glycoproteomics, we prepared a 6plex TMT sample consisting of a labeled yeast peptide mixture spiked with labeled IgM (**Figure 5A**). We labeled IgM peptides separately with individual TMT6 reagents (126-131) and pooled them afterwards in ratios of 10:4:1:1:4:10, respectively. Yeast peptides were labeled with only the first three channels of TMT reagents (126, 127 and 128) and mixed in a 1:1:1 ratio. The labeled IgM peptides were added to the labeled yeast peptides in a 1:1 (w/w) ratio of total peptide amount. We analyzed the mixture by four different MS acquisition strategies: standard MS2 with or without FAIMS, and Glyco-SPS-MS3 with or without FAIMS. All the raw data were processed using the GlycoBinder^5^ pipeline for IgM N-glycopeptide identification and quantification. The predicted ratios of reporter ions of N-glycopeptides between channels are shown in **Figure 5A**. If interference occurs, TMT-labeled and co-isolated yeast peptides may produce additional TMT-reporter ions for the first three channels, 126, 127, 128 (interfered), and may thus hamper the accurate quantification of IgM N-glycopeptides. The ratios of channels 129, 130, and 131 (non-interfered) should not be affected and are expected to remain close to the predicted values.

**Figure 5.**
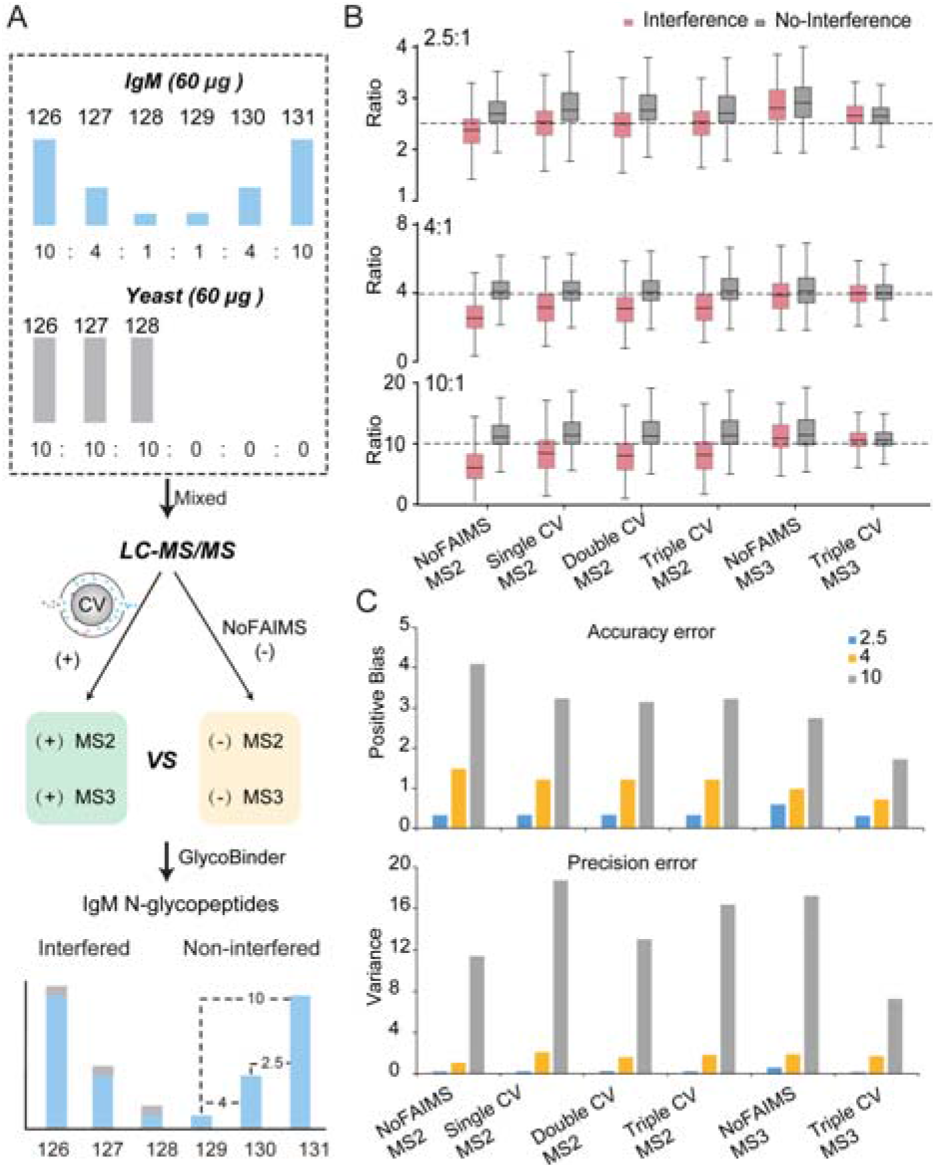
Evaluation of FAIMS for multiplexed quantitative N-glycoproteomics. (A) A scheme of IgM-yeast mixture preparation, MS analysis, and anticipated outcome of interfered N-glycopeptide quantification caused by precursor co-isolation. (B) Ratio distribution of channels with and without yeast interference. Predicted ratios are indicated. Box plots show the median (center line), first and third quartiles (box edges) and 1.5× interquartile range (whiskers). Outliers are not shown. (C) The quantification bias (upper panel) and variance (bottom panel) of each method based on ratios of TMT channels with yeast interference.

In line with our previous observation^5^, the TMT ratios of the interfered channels determined by standard MS2 without FAIMS were skewed, resulting in lowest quantitative accuracy among all the methods (**Figure 5B**). Regarding the analyses using MS2 methods with multiple single-CV FAIMS settings (−40 to −90 V in 5 or 10 V steps; **Figure S9**), we found that the quantitative accuracy was diminished with decreasing CV values. The observed optimum CVs (−45 and −50 V) for quantification are also the optimum CVs for identifying IgM N-glycopeptides (**Figure 1**, see above). Hence, FAIMS can rescue the quantification accuracy to some extent in MS2 analysis of N-glycopeptides. We reasoned that FAIMS suppressed the co-isolation interference by reducing background ions (**Figure S2**). Besides, these optimum CV settings allowed detection/fragmentation of IgM N-glycopeptides with higher MS1 intensities, which were more likely to generate TMT reporter ions with higher signal-to-noise (S/N) ratios and consequently were more resistant to co-isolation interference. The distribution of TMT reporter-ion S/N ratios determined across various single CVs also supports this (**Figure S10**). In the analyses using our Glyco-SPS-MS3 method, we observed, in agreement with our previous results, reduced interference and more accurate TMT ratios compared with the MS2 method (**Figure 5B**). FAIMS-Glyco-SPS-MS3 also improved the accuracy of quantification (**Figure 5B**). Indeed, the combination of FAIMS and Glyco-SPS-MS3 strongly diminished the interference effects and resulted in the accurate determination of TMT ratios of the interfered channels, which matched almost perfectly with the non-interfered ones.

To better access the quantitative accuracy and precision, we calculated the quantification bias (accuracy) and positive variance (precision) of each method (**Figure 5C**, see Methods). We found that the standard MS2 method coupled with FAIMS showed comparable performance for single, double, and triple CV settings in both accuracy and precision. MS2 and MS3 methods with FAIMS lowered the quantitative bias by up to 24% and 49% than that without FAIMS, respectively, increasing the overall quantification accuracy. Interestingly, Glyco-SPS-MS3 without FAIMS led to better accuracy but worse precision than the standard MS2 with and without FAIMS. In contrast to the previous notion^30^ that the MS2 method is less accurate while the SPS-MS3 is less precise, we found that FAIMS in combination with Glyo-SPS-MS3 provided the highest precision and accuracy. Together, our results demonstrate that FAIMS improved multiplexed quantification of site-specific N-glycopeptides in both the MS2 and the Glyco-SPS-MS3 method.

## CONCLUSIONS

In this study, we optimized and thoroughly evaluated the use of FAIMS for multiplexed N-glycoproteomics. We demonstrate that the TMT-labeling changed the FAIMS mobility of N-glycopeptides, which justifies the need for separate optimization of FAIMS CV settings for TMT-labeled and unlabeled N-glycopeptides. Using IgM digest as a model, we show that FAIMS raised the ratio of glycan-oxonium-ion-containing spectra in an LC-MS/MS analysis by up to 76%, increasing the chance of N-glycopeptides being selected for MS/MS and subsequently identified. Compared with the analysis without FAIMS, the optimum CV setting led to a 3.6-fold increase in N-glycopeptide identification from human DG75 cells. TMT labeling and charge states affected FAIMS mobility of IgM N-glycopeptides more significantly than did glycan type, peptide sequence, glycan size and precursor m/z. Lower CVs may cause the neutral loss of glycan moieties, particularly sialic acid, upon FAIMS separation. Finally, we demonstrate that FAIMS further improved the quantification accuracy and precision of both the standard MS2 method and Glyco-SPS-MS3. The combination of Glyco-SPS-MS3 and FAIMS provided the best accuracy and precision and would appear to be the method of choice for multiplexed quantitative N-glycoproteomics.

## ASSOCIATED CONTENT

### Supporting Information

The Supporting Information is available free of charge on the ACS Publications website.

Supplementary figures include comparison of specificity and overlap, the precursor charge state distribution between TMT labeled and unlabeled IgM N-glycopeptides. The influence of charge states and sialic acid on the delta CV. CV scan of synthetic peptides and glycopeptides, comparison of the best single CV for peptides without and with varied glycans, the distribution of precursor mass, precursor m/z, peptide mass and glycan mass for identified TMT labeled and unlabeled IgM N-glycopeptides and DG75 across tested CVs, extract ion chromatography of N-glycopeptides with sialic acid, comparison of accuracy and precision among MS2 methods with single CV, the distribution of signal to noise of reporter ion (PDF).

Supplementary tables include list of all synthetic (glyco)peptides and classification of glycan type and branches. (PDF)

## Author Contributions

K.T.P and H.U. planned and supervised the study. P.F. conceived the study. Y.L.J. contributed to sample preparation. I.S. contributed to data analysis. R.V. contributed to initial measurements with FAIMS pro. T.O. provided materials. P.F., K.T.P. and H.U. wrote the manuscript with help from all other authors.

## Notes

V. R. is an employee of Thermo Fisher Scientific. P.F., Y.L.J., I.S., O.T., K.T.P. and H.U. have no competing interests.

## ACKNOWLEDGMENT

We thank Prof. Daniel Kolarich (Institute for Glycomics, Griffith University, Queensland Australia) for providing the synthetic peptides and glycopeptides. We thank Dr. Wenfeng Zeng (Proteomics and Signal Transduction, Max Planck Institute of Biochemistry, Martinsried, Germany) for pGlyco 2 supports. P.F. was supported by a Manfred Eigen Fellowship from the Max Planck Institute for Biophysical Chemistry. H.U. is funded by a Collaborative Research Center of the Deutsche Forschungsgemeinschaft (SFB1286). K.T.P is supported in part by the LOEWE Center Frankfurt Cancer Institute (FCI) funded by the Hessen State Ministry for Higher Education, Research and the Arts [III L 5-519/03/03.001-(0015)].

