## Supplementary figures for "Evaluation and Optimization of High-Field Asymmetric Waveform Ion Mobility Spectrometry for Multiplexed Quantitative Site-specific N-glycoproteomics"

**Supplementary methods:**

**Preparation of Unlabeled and TMT-labeled Peptides from IgM.** Human IgM purified from human serum (Sigma) was reconstituted at the final concentration of 1 mg/ml in 0.05 M Tris-HCl, 0.2 M sodium chloride, 15 mM sodium azide, pH 8.0. RapiGest SF surfactant (Waters) was dissolved in 50 mM triethylammonium bicarbonate (TEAB) to a concentration of 1% (wt/wt). The prepared RapiGest solution was added to the protein solutions to a final concentration of 0.1% (wt/wt). The proteins were heated at 95 °C for 10 min. After cooling to room temperature (RT), the proteins were reduced and alkylated with 10 mM tris(2-carboxyethyl) phosphine hydrochloride (TCEP, 500 mM stock, Sigma) and 20 mM iodoacetamide at 37 °C for 60 min in the dark. The proteins were digested using trypsin at an enzyme to protein ratio of 1:50 at 37 °C overnight. The digested samples were acidified by adding TFA to a final concentration of 1% (vol/vol). The insoluble particles in the samples were removed by centrifugation at 14,000 × g for 10 min using a benchtop centrifuge. The resulting peptides were cleaned up with an Oasis HBL column and dried in a SpeedVac concentrator. The same digested peptides from IgM were aliquoted into two parts for unlabeled and TMT labeled samples. For TMT labeled IgM peptides, TMT labeling procedure (Thermo Scientific) was performed according to the manufacturer’s instruction. After TMT labeling, the resulting peptides were cleaned up with an Oasis HBL column.

**Cell Culture.** DG75 cells (DSMZ no.: ACC 83) were cultured in RPMI medium (Invitrogen) supplemented with 10–20% (vol/vol) heat-inactivated fetal bovine serum (Invitrogen), penicillin/streptomycin (Invitrogen) and L-glutamine (Invitrogen) at 37 °C with 5% CO_2_. Cells were grown to ~90% confluency before harvesting and provided as cell pellets aliquoted at about 1 × 10^7^ cells per tube.

**Preparation of TMT-labeled Peptides from DG75 Cells.** Cell pellets were lysed in the lysis buffer consisting of 4% SDS (w/v), 50 mM HEPES, pH 8.0. The samples were sonicated for 15 min (15 s on, 15 s off) using Bioruptor at 4 °C. After centrifugation at 14,000 × g for 15 min, the supernatants were collected, and the protein concentration were determined using Pierce BCA Protein Assay Kit (Thermo Scientific). The proteins were reduced and alkylated with 10 mM TCEP (500 mM stock, Sigma) and 20 mM iodoacetamide at 37 °C for 60 min in the dark. A mixture of magnetic beads, including Sera-Mag SpeedBeads with a hydrophilic surface (GE Healthcare, cat.no. 45152101010250, Magnetic Carboxylate Modified), and Sera-Mag SpeedBeads with a hydrophobic surface (GE Healthcare, cat.no. 65152105050250, Magnetic Carboxylate Modified) was rinsed twice with water on a magnetic rack prior to use. The beads were added to protein lysates at the optimal working ratio of 10:1 (wt/wt, beads to proteins). The required minimum bead concentration is 0.5 μg/μL in order to provide sufficient surface for the immobilization of aggregated proteins. We then added acetonitrile (ACN) to protein lysates to a final percentage of 70% (vol/vol). The samples were allowed to stay off the rack for 10 min at room temperature (RT), followed by resting on the magnetic rack for 2 min at RT. The supernatant was discarded, and the beads were then washed for three times with 80% (vol/vol) ethanol. Beads were resuspended in 50 mM TEAB containing sequencing grade modified trypsin (1:50 of enzyme to protein amount) and incubated at 37 °C for 4 h or overnight in a ThermoMixer with mixing at 800 rpm. After digestion, we placed the tubes on a magnetic rack for a few minutes and transferred the supernatant to a fresh tube. TMT labeling procedure (Thermo Scientific) was performed according to the manufacturer’s instruction.

**Enrichment of Glycopeptides from DG75 Samples using Zwitterionic Hydrophilic Interaction Liquid Chromatography (ZIC-HILIC).** Digested peptides from DG75 were acidified by adding 10% (vol/vol) trifluoroacetic acid (TFA) to a final concentration of 1% followed by centrifugation at 14,000 × g for 20 min. The supernatant was dried in a SpeedVac concentrator. We re-dissolved the dried peptides in loading buffer consisting of 80% (vol/vol) ACN and 1% TFA and maintained the peptide concentration at around 4 mg/ml. Meanwhile, we weighted out the ZIC-HILIC beads (5 μm, Welch) according to the peptide to bead ratio of 1:50 (wt/wt) and washed three times using the loading buffer. We loaded all beads onto a 200 μL pipette tip pre-packed with coffee filter. Samples were loaded five times (at least 4 min each time) followed by three times wash with loading buffer. The retained glycopeptides were eluted with 100 μL 0.1% TFA twice. The collected eluates were dried in a SpeedVac concentrator.

**IgM and Yeast Peptide Interference Model**. Yeast protein prepared from S. cerevisiae cells were purchased from Promega (Cat. V7341). Yeast proteins in 6.5 M urea/50 mM Tris-HCl (pH 8) at a protein concentration of 10 mg/ml were thawed on ice. Protein reduction and alkylation were the same as mentioned above. After diluting urea to 1 M using 50 mM Tris-HCl (pH 8), trypsin (Promega) was added at a trypsin:protein ratio of 1:50 at 37 °C. After overnight incubation, the samples were acidified by adding TFA to a final concentration of 1% (vol/vol). The insoluble particles in the samples were removed by centrifuging at 14,000 × g for 10 min using a benchtop centrifuge. The samples were cleaned up with Oasis HBL columns. The resulted peptides were dried in a SpeedVac concentrator.

We separately labeled IgM digests with individual TMT6 reagents and pooled them together afterward with the ratio of 10:4:1:1:4:10. In contrast, yeast peptides were labeled with only the first three channels of TMT6 reagents (126, 127, and 128) and mixed equally (**Fig. 5A**). We then spiked the pooled IgM peptides into the yeast peptide mixture with an equal amount. The mixed samples were then cleaned up with Oasis HBL columns followed by analysis with either Glyo-SPS-MS3 or standard MS2 methods (see below).

**LC-MS/MS Analysis.** Peptides were resuspended in 5% ACN, 0.1% FA and subjected for LC-MS/MS analysis using Orbitrap Exploris 480 or Orbitrap Fusion Tribrid Mass Spectrometers (Thermo Fisher Scientific), both coupled to a Dionex UltiMate 3000 UHPLC system (Thermo Fisher Scientific). Unless noted otherwise, all samples were analyzed using a C18 trap column (3 cm long; inner diameter, 100 μm; outer diameter, 360 μm) and a home-made analytical column (ReproSil-Pur 120 C18-AQ, 1.9 µm pore size, 75 µm inner diameter, Dr. Maisch GmbH, 30 cm), the gradient at a flow rate of 300 nl/min, mobile phase A and B consisting of 0.1% (vol/vol) formic acid (FA) and 80% ACN, 0.08% FA, respectively. The one-hour gradient started at 10% B at 3 min, increased to 45% B at 47 min, and then to 90% B in 0.1 min. After washing with 90% B for 5 min, the column was re-equilibrated with 5% B. The two hours gradient started at 10% B at 3 min, increased to 45% B at 107 min, and then to 90% B in 0.1 min. After washing with 90% B for 5 min, the column was re-equilibrated with 5% B. The three hours gradient started at 10% B at 3 min, increased to 45% B at 166 min, and then to 90% B in 0.1 min. After washing with 90% B for 5 min, the column was re-equilibrated with 5% B.

The MS settings for the Glyco-SPS-MS3 method measured on Orbitrap Fusion are the same with previous publication^1^. For standard MS2 method on Orbitrap Exploris 480, the following optimized MS settings were used. MS1 settings: Orbitrap Resolution-120 k, Scan Range-350–2,000, Maximum injection time-50 ms, AGC target-300% (3e6), RF Lens-40%, and Data Type-Profile. MS2 settings: Precursor selection range-700-2000 m/z, Isolation - window-1.6 m/z, Scan range mode-Auto normal, First mass-120, Normalized collision energy (%)-30/35/40, Detector type-Orbitrap, Orbitrap resolution-15 K, Maximum injection time-100 ms, AGC target-(50%)5e4, and Data type-Profile.

**Data Analysis.** For intact glycopeptide identification and quantification, .raw files were processed via GlycoBinder^1^. Parameters used for pGlyco 2 include fully specific trypsin digestion with maximal two missed cleavage and mass tolerance for precursors and fragment ions of 10 and 20 ppm, respectively. We considered cysteine carbamidomethylation and TMT0 (or TMT6) on peptide N-termini and lysine residues as fixed modifications and methionine oxidation as a variable modification. The reviewed human protein database was downloaded from Swiss-Prot (March 2018, human, 20,303 entries). For the identification of IgM glycopeptides, we included only the sequences of human IGHM and IGJ in the FASTA file. Only GPSMs with PepScore ≥ 7 and GlyScore ≥ 8 reported by pGlyco 2 were used for following analysis. For the DG75 samples, we used the total FDR ≤ 2% for both the first and second database search.

For the IgM-Yeast interference model, we concatenated a reviewed S.cerevisiae protein database (Uniprot, 4,525 entries, March 2020, strain ATCC 204508 / S288c) with the sequences of human IGHM and IGJ protein for database search. FDR < 2% for both the first and second database search was used. A global analysis of all TMT ion ratios obtained for yeast and human peptides is shown for LC-MS/MS analyses performed with and without FAIMS. GPSMs of IgM glycopeptides that did not show reporter ion intensities in all TMT channels were filtered out.

**Classification of Glycan Compositions.** Glycan compositions were classified as (1) “oligomannose” if they consist of two HexNAc and more than five Hex; (2) “paucimannose” if they match the composition of Hex(1-4)Fuc(0-1)HexcNAc(2), where the number of HexNAc is 2, the number of Fuc is 0 or 1, and the number of Hex is 1 to 4; (3) “short” if they match the composition of Hex(0-2)Fuc(0-1)HexNAc(1-3); (4) “complex” if they contain Hex-HexNAc antenna(e) attached to the common N-glycan core (Hex3HexNAc2); and (5) “hybrid” if they contain both Hex-HexNAc antenna(e) and extra Hex residues attached to the N-glycan core. There is no general composition for complex and hybrid types, as their structures are relatively more complex. We additionally include a “complex/hybrid” type for the glycan compositions that cannot be assigned as complex or hybrid types unambiguously.

We propose the potential number of branches of each glycan composition based on the glycan types. The number of branches in complex and hybrid type glycans was determined by the lowest possible number of antennae extended from the N-glycan core based on our understanding of human N-glycosylation biosynthesis. For the oligomannose type, the number of branches is 3. For the paucimannose and short glycan types, the number of branches is 1 or 2. Please see **Table S2** for the complete list of glycan compositions and their classifications.

**Supplementary Figures:**

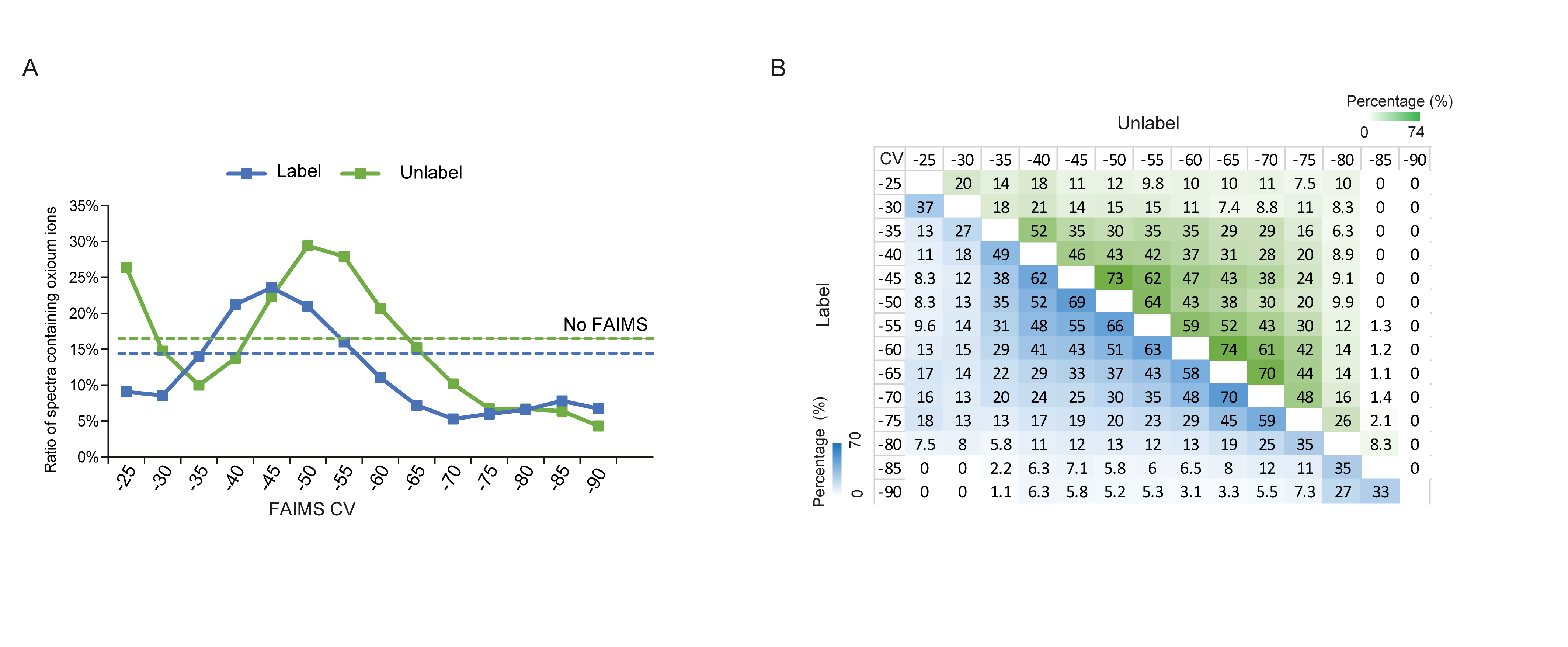

**Figure S1**. **The effects of FAIMS CV settings on glycopeptide identification.** (A) Ratio of MS2 spectra containing 204.078 to total MS2 scans in a single LC-MS/MS run using varying CVs. Dashed lines represent the analyses without FAIMS. (B) Overlaps of unique TMT-labeled (blue) and unlabeled glycopeptides identified in LC-MS/MS runs using different CVs.

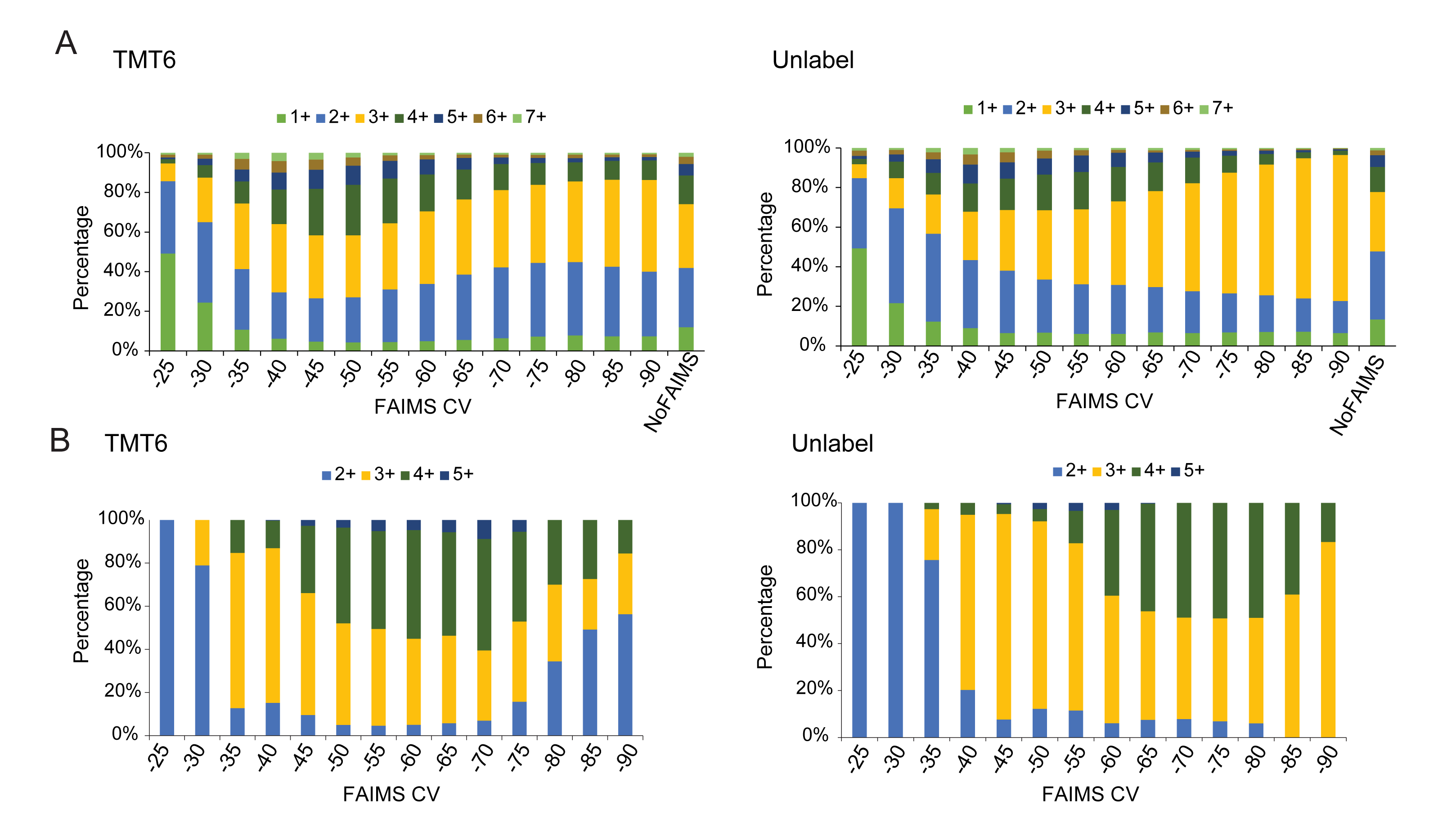

**Figure S2. Distributions of precursor charge state detected in an LC-MS/MS measurement using varying FAIMS CVs.** (A) Charge state distribution of all MS1 features detected from TMT-labeled (left) or unlabeled (right) IgM samples across tested CVs. (B) Charge state distribution of identified glycopeptides with or without TMT-labeling across tested CVs.

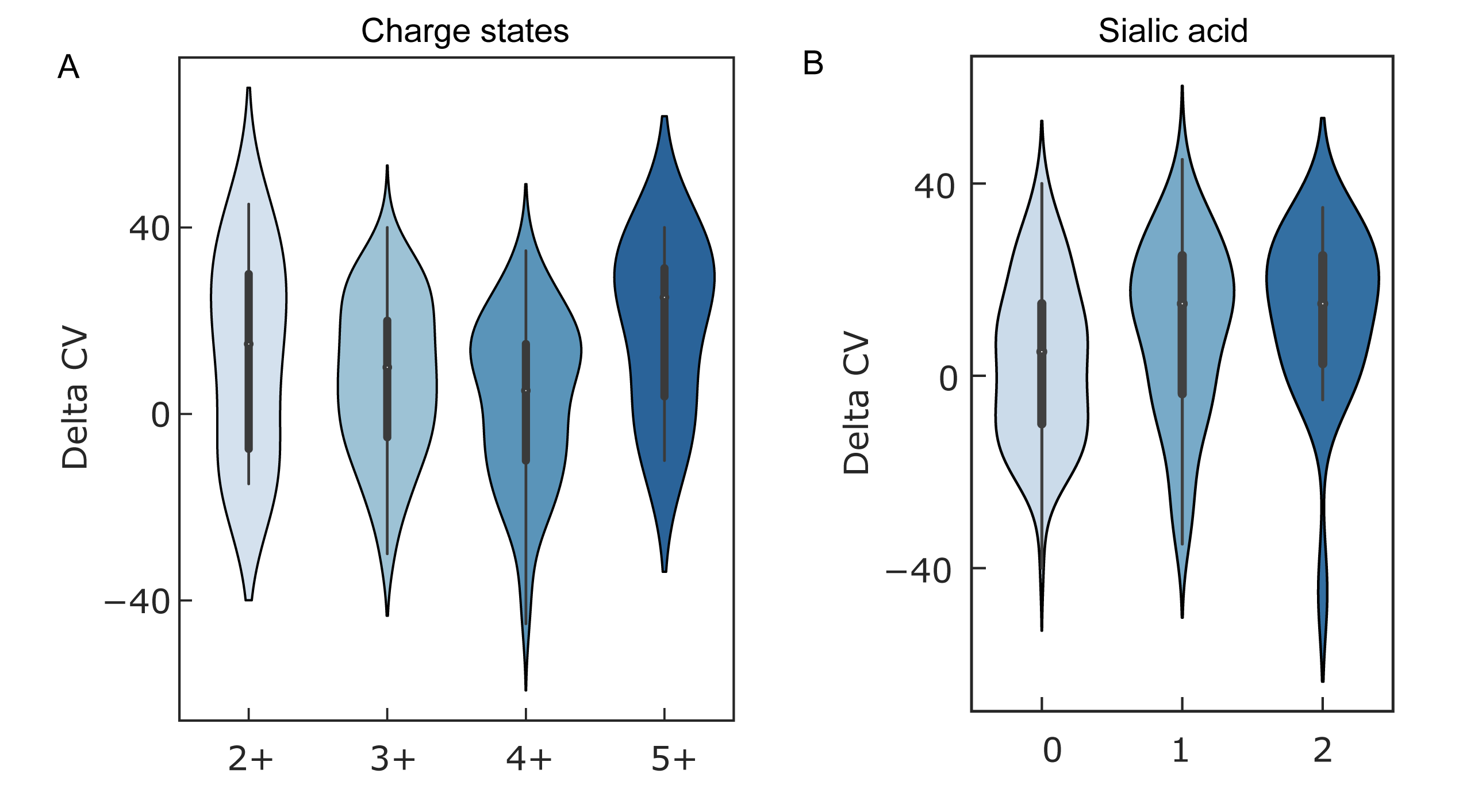

**Figure S3. The influence of charge states and the number of sialic acid on the delta CV.** We detected differences (i.e., delta CVs) in the optimal CVs between TMT-labeled glycopeptides and their unlabeled counterparts. (A) The distribution of delta CVs for N-glycopeptides with different charge states. (B) The distribution of delta CV for N-glycopeptides with different numbers of sialic acids.

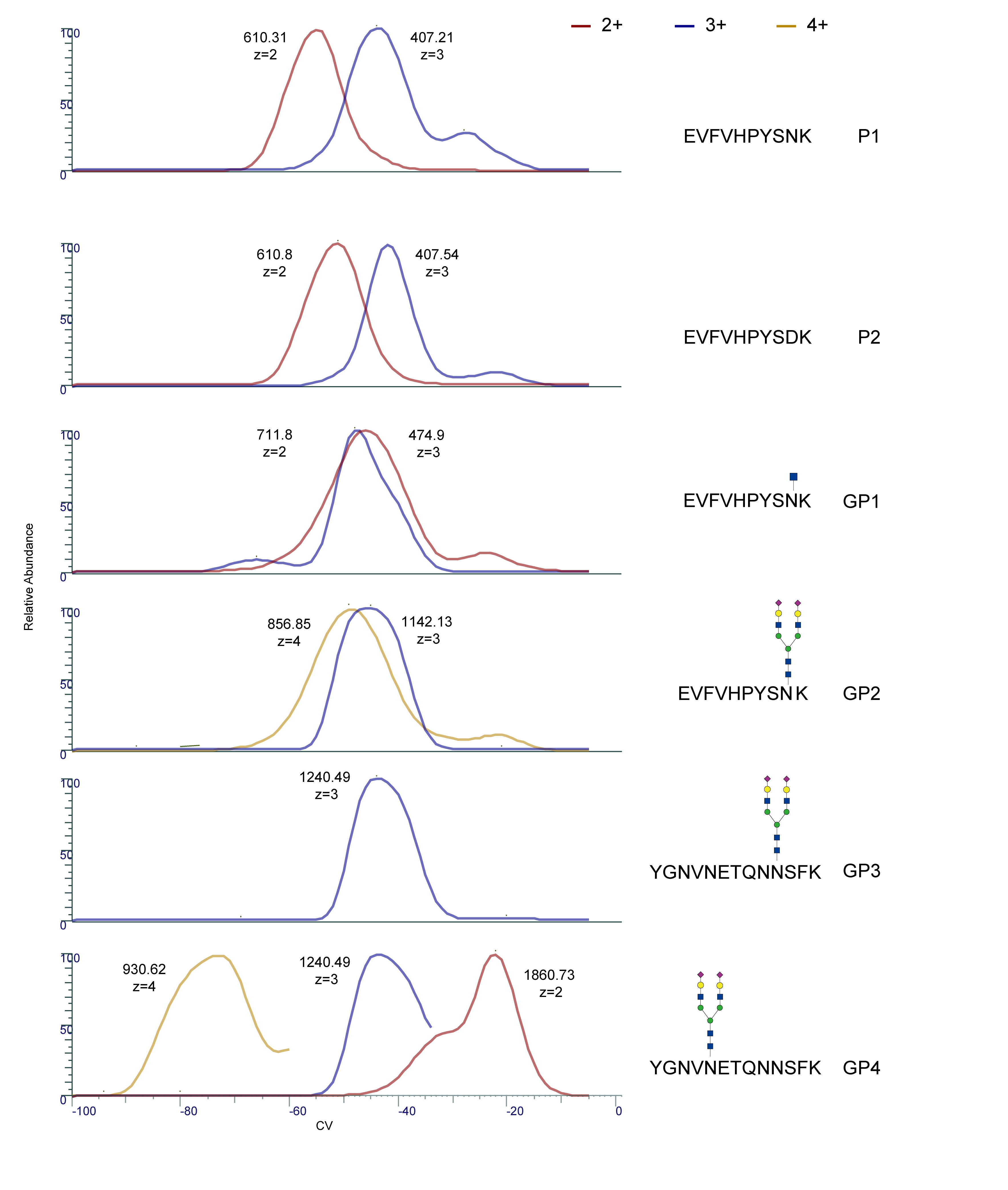

**Figure S4. CV scans of synthetic peptides and glycopeptides from -100 V to 0 V (in 1 V step).** Ion signals of all (glyco)peptides with different charge states detected with varying CVs were extracted and shown in different colors. The m/z used for extracting signals are marked.

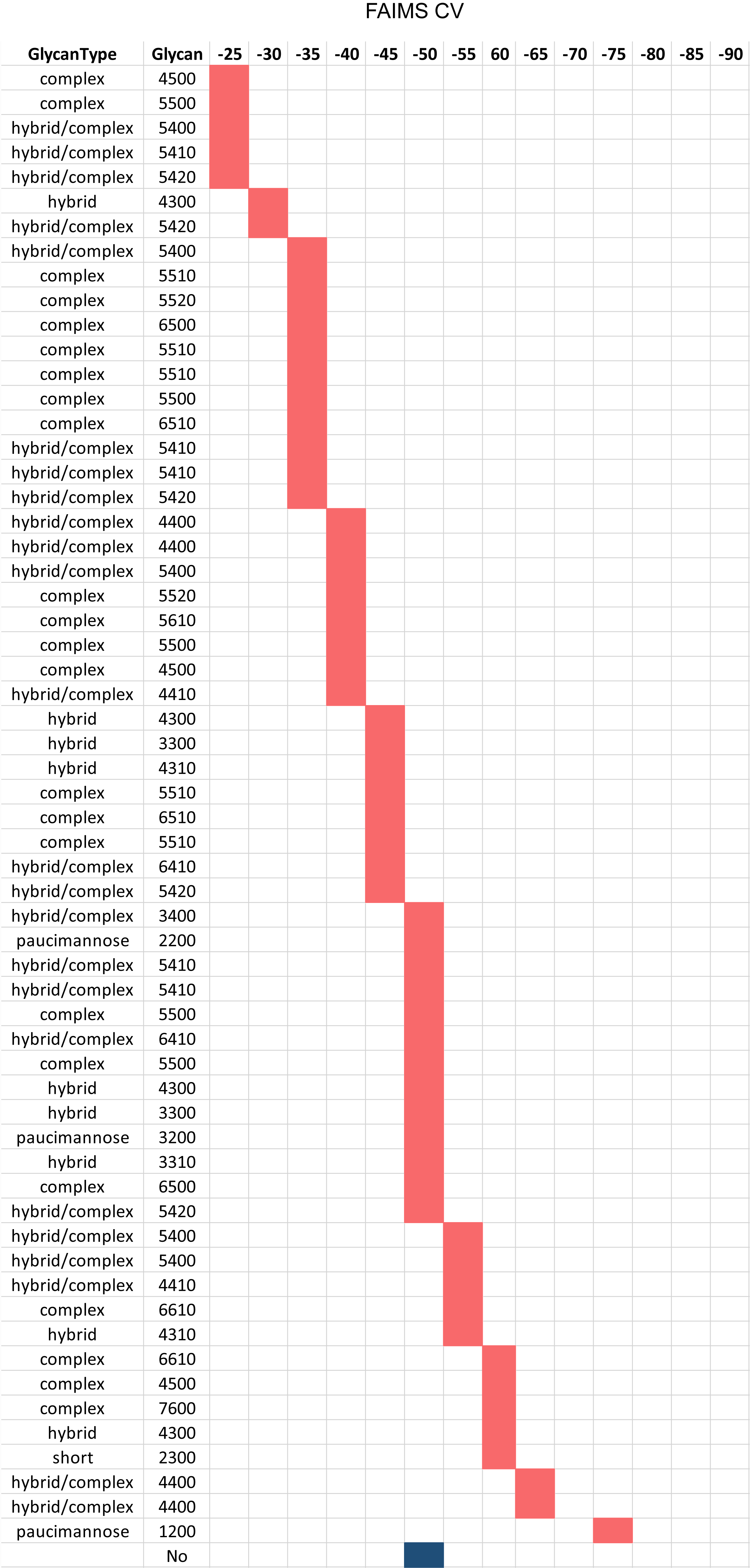

**Figure S5. Comparison of the best single CV for peptides without (blue) and with glycans (red).** We selected the (glyco)peptides sharing a common peptide sequence of E**N**ISDPTSPLR for comparison. The best CV is the CV that provided the maximum intensity of each (glyco)peptide among all tested CVs. The glycan compositions are represented by the number of Hex, HexNAc, NeuAc, and Fuc. Each glycan composition was classified into different glycan types according to the criteria described in the Method section.

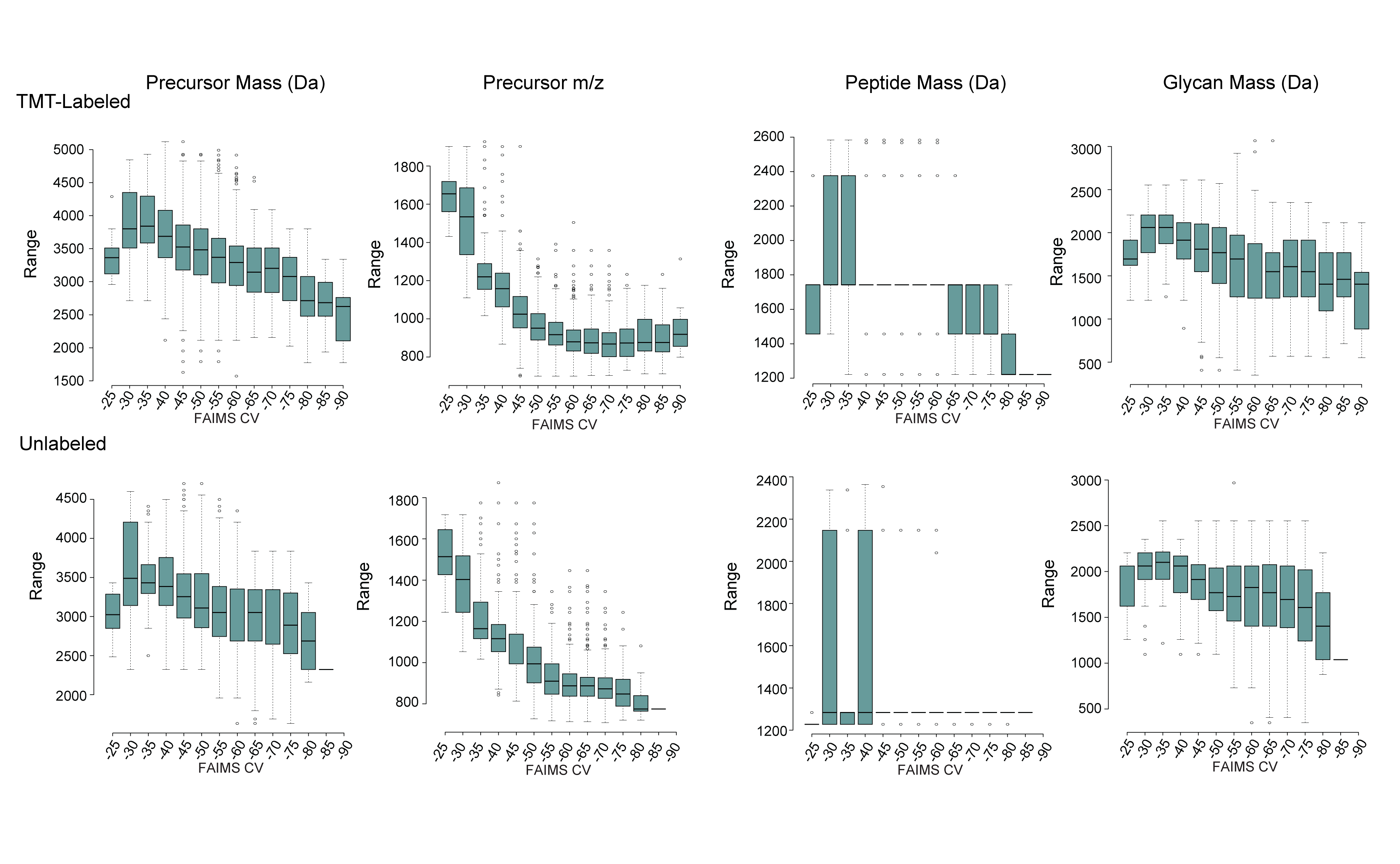

**Figure S6**. **The distributions of precursor mass, precursor m/z, peptide mass, and glycan mass for identified TMT-labeled and unlabeled IgM glycopeptides across the tested CVs.** Boxplots show the median (centerline), first and third quartiles (box edges) and 1.5 × the interquartile range (whiskers).

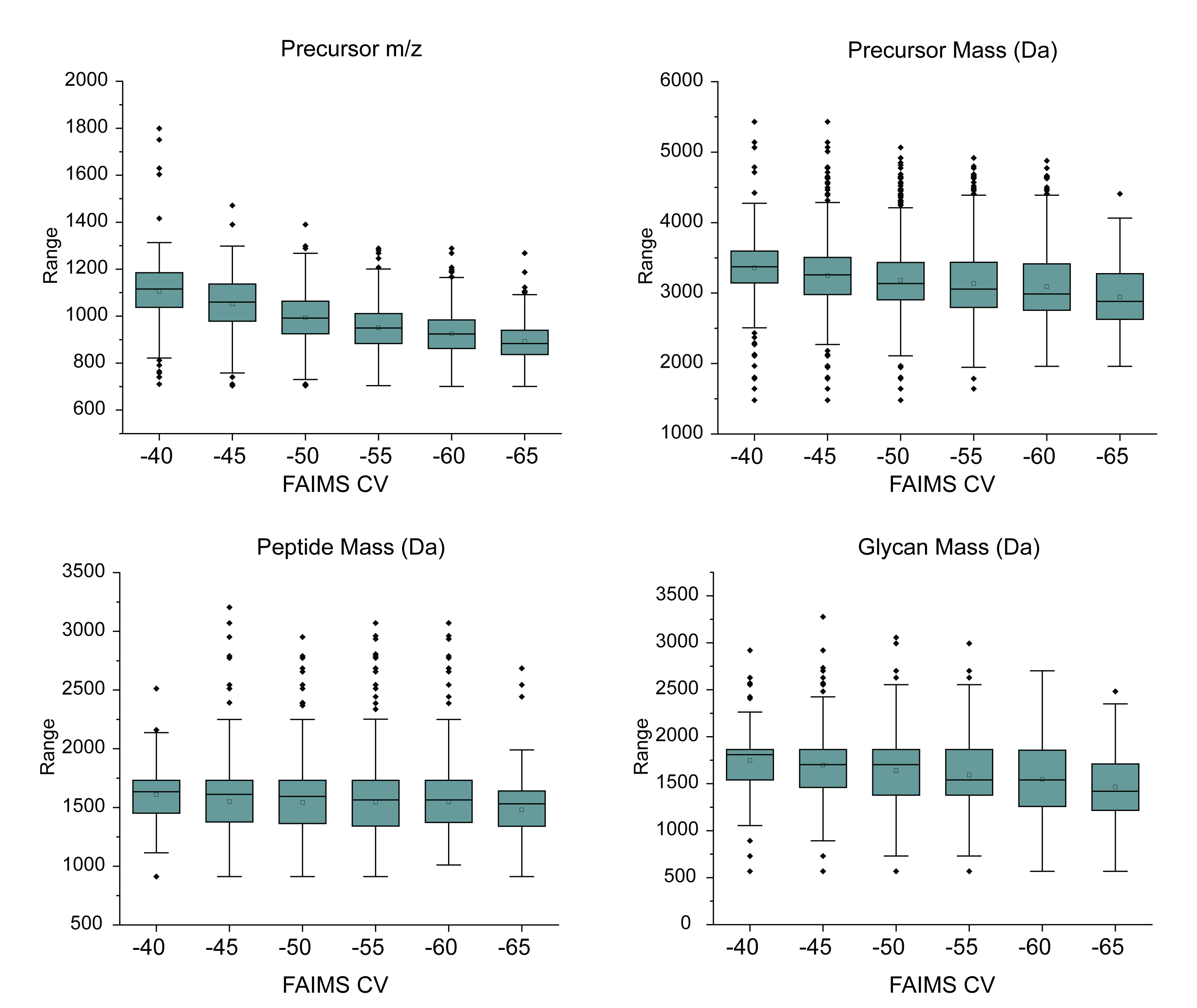

**Figure S7**. **The distributions of precursor m/z, precursor mass, glycan mass, and peptide mass of glycopeptides identified in DG75 across tested CVs.** Boxplots show the median (centerline), first and third quartiles (box edges) and 1.5 × the interquartile range (whiskers).

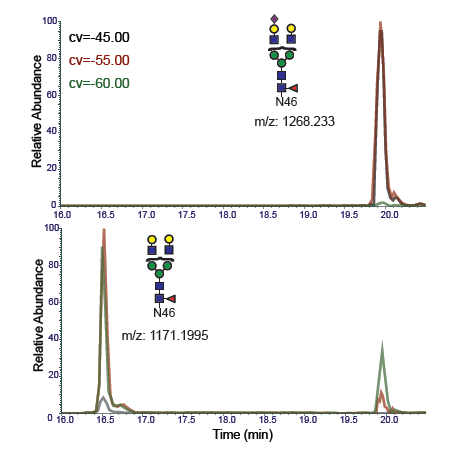

**Figure S8. Neutral loss of NeuAC moiety upon FAIMS separation.** Extracted ion chromatograms (XIC) of the two-antenna glycopeptides with or without terminal NeuAC were overlapped. XICs from different CVs were shown in different colors.

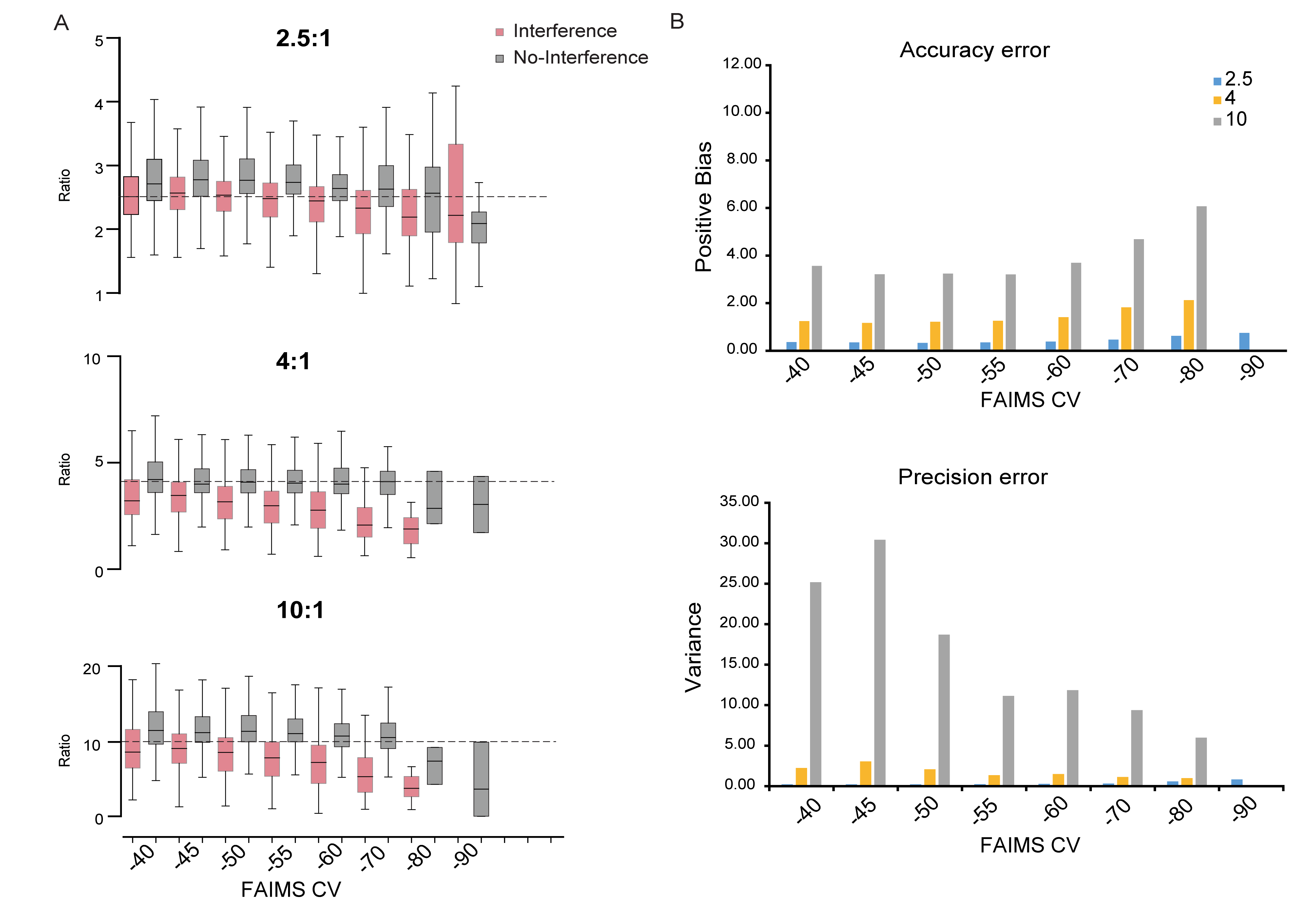

**Figure S9**. **Effects of FAIMS on the accuracy and precision of multiplexed quantitative N-glycoproteomics.** The IgM-yeast mixture was analyzed by LC-FAIMS-MS2 using varied single CVs. TMT reporter ratios were determined using GlycoBinder pipeline. (A) Ratio distribution of channels with (red) and without (grey) yeast interference. Predicted ratios are indicated. Boxplots show the median (centerline), first and third quartiles (box edges) and 1.5× the interquartile range (whiskers). Outliers are not shown. (B) Quantification bias (upper panel) and variance (bottom panel) of each measurement based on ratios of TMT channels with yeast interference.

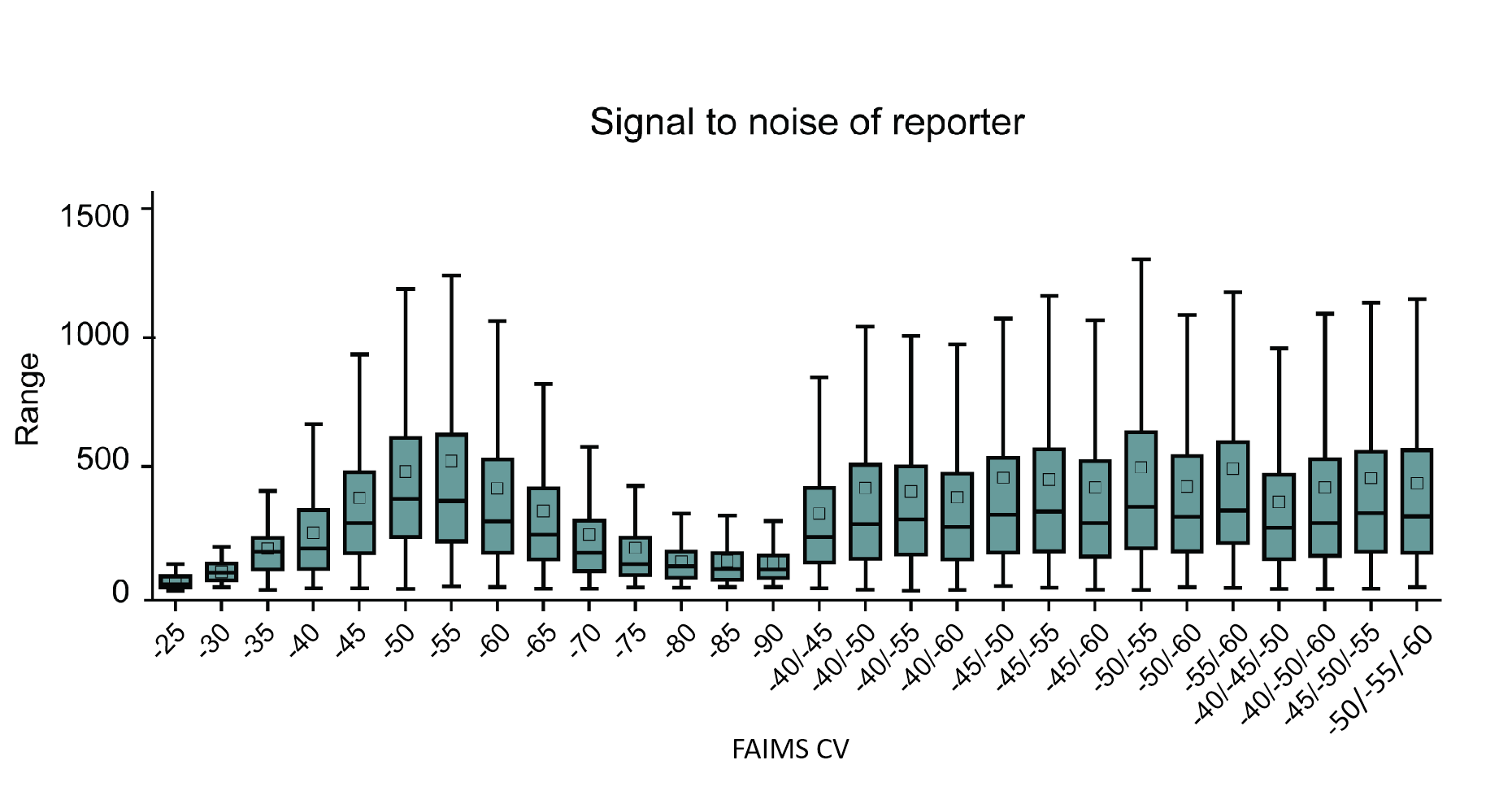

**Figure S10**. **The distribution of signal to noise of reporter ion across tested CV settings.** Boxplots show the median (centerline), mean (squares), first and third quartiles (box edges) and 1.5 × the interquartile range (whiskers). Outliers are not shown.

**Supplementary Tables:**

**Table S1: List of all synthetic (glyco)peptides and their detected optimal CVs.**

|  | **Peptide sequences** | **Modification** | **Charge** | **Monoisotopic m/z** | **Peak CV (v)** |
| --- | --- | --- | --- | --- | --- |
| P1 | EVFVHPYS**N**K | - | 2+ | 610.31 | -55 |
|  |  |  | 3+ | 407.21 | -44 |
| P2 | EVFVHPYS**D**K | - | 2+ | 610.8009 | -53 |
|  |  |  | 3+ | 407.54 | -42 |
| GP1 | EVFVHPYSN^*^K | GlcNAc | 2+ | 711.8 | -46 |
|  |  |  | 3+ | 474.9 | -47 |
| GP2 | EVFVHPYSN^*^K | 5Hex4HexNAc2NeuAc | 3+ | 1142.13 | -45 |
|  |  |  | 4+ | 856.85 | -49 |
| GP3 | YGNVNETQNN^*^SFK |  | 3+ | 1240.49 | -46 |
| GP4 | YGNVN^*^ETQNNSFK |  | 2+ | 1860.73 | -23 |
|  |  |  | 3+ | 1240.49 | -44 |
|  |  |  | 4+ | 930.62 | -70 |

Note: The amino acid difference between P1 and P2 is marked in red. The glycosylation sites are marked using a red star.

**Table S2. The putative glycan types and branches.**

| Number of glycan units | | | |  |  |
| --- | --- | --- | --- | --- | --- |
| Hex | HexNAc | Fucose | NeuAc | glycan.Type | number.of.branches |
| 0 | 1 | 1 | 0 | short | 1 |
| 0 | 2 | 0 | 0 | short | 1 |
| 0 | 2 | 1 | 0 | short | 1 |
| 1 | 2 | 0 | 0 | paucimannose | 1 |
| 1 | 2 | 1 | 0 | paucimannose | 1 |
| 2 | 2 | 0 | 0 | paucimannose | 1 |
| 2 | 2 | 1 | 0 | paucimannose | 1 |
| 3 | 2 | 0 | 0 | paucimannose | 2 |
| 3 | 2 | 1 | 0 | paucimannose | 2 |
| 4 | 2 | 0 | 0 | paucimannose | 2 |
| 4 | 2 | 1 | 0 | paucimannose | 2 |
| 5 | 2 | 0 | 0 | oligo-mannose | 3 |
| 5 | 2 | 1 | 0 | oligo-mannose | 3 |
| 6 | 2 | 0 | 0 | oligo-mannose | 3 |
| 7 | 2 | 0 | 0 | oligo-mannose | 3 |
| 8 | 2 | 0 | 0 | oligo-mannose | 3 |
| 8 | 2 | 1 | 0 | oligo-mannose | 3 |
| 9 | 2 | 0 | 0 | oligo-mannose | 3 |
| 9 | 2 | 1 | 0 | oligo-mannose | 3 |
| 10 | 2 | 0 | 0 | oligo-mannose | 3 |
| 11 | 2 | 0 | 0 | oligo-mannose | 3 |
| 12 | 2 | 0 | 0 | oligo-mannose | 3 |
| 1 | 3 | 0 | 0 | short | 1 |
| 1 | 3 | 1 | 0 | short | 1 |
| 2 | 3 | 0 | 0 | short | 2 |
| 2 | 3 | 1 | 0 | short | 2 |
| 3 | 3 | 0 | 0 | hybrid | 2 |
| 3 | 3 | 0 | 1 | hybrid | 2 |
| 3 | 3 | 1 | 0 | hybrid | 2 |
| 3 | 3 | 1 | 1 | hybrid | 2 |
| 3 | 3 | 2 | 0 | hybrid | 2 |
| 4 | 3 | 0 | 0 | hybrid | 2 |
| 4 | 3 | 0 | 1 | hybrid | 2 |
| 4 | 3 | 1 | 0 | hybrid | 2 |
| 4 | 3 | 1 | 1 | hybrid | 2 |
| 4 | 3 | 1 | 2 | hybrid | 2 |
| 4 | 3 | 2 | 0 | hybrid | 2 |
| 4 | 3 | 2 | 2 | hybrid | 2 |
| 4 | 3 | 3 | 0 | hybrid | 2 |
| 5 | 3 | 0 | 0 | hybrid | 3 |
| 5 | 3 | 0 | 1 | hybrid | 3 |
| 5 | 3 | 1 | 0 | hybrid | 3 |
| 5 | 3 | 1 | 1 | hybrid | 3 |
| 5 | 3 | 2 | 0 | hybrid | 3 |
| 5 | 3 | 2 | 2 | hybrid | 3 |
| 5 | 3 | 3 | 0 | hybrid | 3 |
| 6 | 3 | 0 | 0 | hybrid | 3 |
| 6 | 3 | 0 | 1 | hybrid | 3 |
| 6 | 3 | 0 | 2 | hybrid | 3 |
| 6 | 3 | 1 | 0 | hybrid | 3 |
| 6 | 3 | 1 | 1 | hybrid | 3 |
| 6 | 3 | 1 | 2 | hybrid | 3 |
| 6 | 3 | 2 | 1 | hybrid | 3 |
| 6 | 3 | 3 | 0 | hybrid | 3 |
| 7 | 3 | 0 | 0 | hybrid | 3 |
| 8 | 3 | 0 | 0 | hybrid | 3 |
| 2 | 4 | 0 | 0 | hybrid/complex | 2 |
| 2 | 4 | 1 | 0 | hybrid/complex | 2 |
| 3 | 4 | 0 | 0 | hybrid/complex | 2/3 |
| 3 | 4 | 0 | 1 | hybrid/complex | 2/3 |
| 3 | 4 | 1 | 0 | hybrid/complex | 2/3 |
| 3 | 4 | 1 | 1 | complex | 2 |
| 3 | 4 | 3 | 0 | complex | 2 |
| 4 | 4 | 0 | 0 | hybrid/complex | 2/3 |
| 4 | 4 | 0 | 1 | hybrid/complex | 2/3 |
| 4 | 4 | 0 | 2 | hybrid/complex | 2/3 |
| 4 | 4 | 1 | 0 | hybrid/complex | 2/3 |
| 4 | 4 | 1 | 1 | hybrid/complex | 2/3 |
| 4 | 4 | 2 | 0 | hybrid/complex | 2/3 |
| 4 | 4 | 2 | 1 | hybrid/complex | 2/3 |
| 4 | 4 | 3 | 0 | hybrid/complex | 2/3 |
| 4 | 4 | 3 | 1 | hybrid/complex | 2/3 |
| 4 | 4 | 4 | 0 | hybrid/complex | 2/3 |
| 5 | 4 | 0 | 0 | hybrid/complex | 2/4 |
| 5 | 4 | 0 | 1 | complex | 2 |
| 5 | 4 | 0 | 2 | complex | 2 |
| 5 | 4 | 0 | 3 | complex | 2 |
| 5 | 4 | 1 | 0 | hybrid/complex | 2/4 |
| 5 | 4 | 1 | 1 | complex | 2 |
| 5 | 4 | 1 | 2 | complex | 2 |
| 5 | 4 | 2 | 0 | hybrid/complex | 2/4 |
| 5 | 4 | 2 | 1 | complex | 2 |
| 5 | 4 | 2 | 2 | complex | 2 |
| 5 | 4 | 3 | 0 | hybrid/complex | 2/4 |
| 5 | 4 | 3 | 1 | complex | 2 |
| 5 | 4 | 4 | 0 | complex | 2 |
| 5 | 4 | 5 | 0 | complex | 2 |
| 6 | 4 | 0 | 0 | hybrid/complex | 2/4 |
| 6 | 4 | 0 | 1 | hybrid/complex | 2/4 |
| 6 | 4 | 1 | 0 | hybrid/complex | 2/4 |
| 6 | 4 | 1 | 1 | hybrid/complex | 2/4 |
| 6 | 4 | 2 | 0 | hybrid/complex | 2/4 |
| 6 | 4 | 3 | 0 | hybrid/complex | 2/4 |
| 7 | 4 | 0 | 0 | hybrid/complex | 2/4 |
| 7 | 4 | 0 | 1 | hybrid/complex | 2/4 |
| 7 | 4 | 1 | 0 | hybrid/complex | 2/4 |
| 7 | 4 | 1 | 1 | hybrid/complex | 2/4 |
| 7 | 4 | 2 | 0 | hybrid/complex | 2/4 |
| 3 | 5 | 0 | 0 | complex | 2/3 |
| 3 | 5 | 1 | 0 | complex | 2/3 |
| 3 | 5 | 3 | 1 | complex | 2/3 |
| 3 | 5 | 4 | 0 | complex | 2/3 |
| 4 | 5 | 0 | 0 | complex | 2/3 |
| 4 | 5 | 0 | 1 | complex | 2/3 |
| 4 | 5 | 0 | 2 | complex | 2/3 |
| 4 | 5 | 1 | 0 | complex | 2/3 |
| 4 | 5 | 1 | 1 | complex | 2/3 |
| 4 | 5 | 2 | 0 | complex | 2/3 |
| 4 | 5 | 3 | 0 | complex | 2/3 |
| 4 | 5 | 4 | 0 | complex | 2/3 |
| 5 | 5 | 0 | 0 | complex | 2/3 |
| 5 | 5 | 0 | 1 | complex | 2/3 |
| 5 | 5 | 0 | 2 | complex | 2/3 |
| 5 | 5 | 1 | 0 | complex | 2/3 |
| 5 | 5 | 1 | 1 | complex | 2/3 |
| 5 | 5 | 1 | 2 | complex | 2/3 |
| 5 | 5 | 2 | 0 | complex | 2/3 |
| 5 | 5 | 2 | 1 | complex | 2/3 |
| 5 | 5 | 3 | 0 | complex | 2/3 |
| 5 | 5 | 3 | 1 | complex | 2/3 |
| 5 | 5 | 5 | 0 | complex | 2/3 |
| 6 | 5 | 0 | 0 | complex | 2/3 |
| 6 | 5 | 0 | 1 | complex | 2/3 |
| 6 | 5 | 0 | 2 | complex | 2/3 |
| 6 | 5 | 1 | 0 | complex | 2/3 |
| 6 | 5 | 1 | 1 | complex | 2/3 |
| 6 | 5 | 1 | 2 | complex | 2/3 |
| 6 | 5 | 2 | 0 | complex | 2/3 |
| 6 | 5 | 2 | 1 | complex | 2/3 |
| 6 | 5 | 3 | 0 | complex | 2/3 |
| 6 | 5 | 4 | 3 | complex | 2/3 |
| 7 | 5 | 1 | 0 | complex | 2/3 |
| 3 | 6 | 2 | 0 | complex | 2/4 |
| 3 | 6 | 3 | 0 | complex | 2/4 |
| 3 | 6 | 4 | 0 | complex | 2/4 |
| 4 | 6 | 0 | 0 | complex | 4 |
| 4 | 6 | 0 | 1 | complex | 4 |
| 4 | 6 | 1 | 0 | complex | 4 |
| 5 | 6 | 0 | 0 | complex | 3/4 |
| 5 | 6 | 0 | 1 | complex | 3/4 |
| 5 | 6 | 0 | 2 | complex | 3/4 |
| 5 | 6 | 1 | 0 | complex | 3/4 |
| 5 | 6 | 1 | 1 | complex | 3/4 |
| 5 | 6 | 2 | 0 | complex | 3/4 |
| 5 | 6 | 2 | 1 | complex | 3/4 |
| 5 | 6 | 3 | 1 | complex | 3/4 |
| 5 | 6 | 5 | 0 | complex | 3/4 |
| 6 | 6 | 0 | 0 | complex | 3/4 |
| 6 | 6 | 0 | 1 | complex | 3/4 |
| 6 | 6 | 2 | 2 | complex | 3/4 |
| 6 | 6 | 3 | 1 | complex | 3/4 |
| 6 | 6 | 4 | 1 | complex | 3/4 |
| 7 | 6 | 4 | 0 | complex | 2/3/4 |
| 9 | 6 | 0 | 1 | complex | 3/4 |
| 10 | 6 | 1 | 1 | complex | 4 |
| 10 | 6 | 2 | 1 | complex | 4 |
| 3 | 7 | 2 | 0 | complex | 4/5 |
| 3 | 7 | 4 | 0 | complex | 4/5 |
| 4 | 7 | 1 | 1 | complex | 4/5 |
| 4 | 7 | 3 | 0 | complex | 4/5 |
| 5 | 7 | 1 | 1 | complex | 4/5 |
| 7 | 7 | 2 | 1 | complex | 5 |
| 8 | 7 | 1 | 2 | complex | 5 |
| 10 | 7 | 0 | 0 | complex | 5 |
| 11 | 7 | 1 | 0 | complex | 5 |
| 3 | 8 | 0 | 0 | complex | 5 |
| 10 | 8 | 3 | 1 | complex | 5 |
